# Smelling in the dark: phylogenomic insights on the chemosensory system of a subterranean beetle

**DOI:** 10.1101/2020.10.22.350173

**Authors:** Pau Balart-García, Alexandra Cieslak, Paula Escuer, Julio Rozas, Ignacio Ribera, Rosa Fernández

**Author notes:** Senior authors (Ignacio Ribera, Rosa Fernández).

## Abstract

The chemosensory system has experienced relevant changes in subterranean animals, facilitating the orientation into darkness via the perception of specific chemical signals critical to survive in this particular environment. However, the genomic basis of chemoreception in cave-dwelling fauna is largely unexplored. We generated *de novo* transcriptomes for antennae and body samples of the troglobitic beetle *Speonomus longicornis* (whose characters suggest an extreme adaptation to the deep subterranean) in order to interrogate the evolutionary origin and diversification of the chemosensory gene repertoire across coleopterans through a phylogenomic approach. Our results suggested a diminished diversity of odorant and gustatory gene repertoires compared to polyphagous epigean beetles. Moreover, *S. longicornis* showed a large diversity of odorant-binding proteins, suggesting an important role of these proteins in capturing airborne chemical cues. We identified a gene duplication in the ionotropic co-receptor I R25a, a highly conserved single-copy gene in protostomes involved in thermal and humidity sensing. In addition, no homologous genes to sugar receptors or the ionotropic receptor IR41a were detected. Our findings suggest that the chemosensory gene repertoire of this cave beetle may have been reshaped by the low complexity of chemical signals of this particular environment, and that gene duplication and loss may have played an important role in the evolution of genes involved in chemoreception. Altogether, our results shed light on the genomic basis of chemoreception in a cave-dwelling invertebrate and pave the road towards understanding the genomic underpinnings of adaptation to the subterranean lifestyle at a deeper level.

## INTRODUCTION

Major lifestyle transitions in insects, such as the conquest of terrestrial habitats, the flight or host plant interactions, are often followed by dramatic shifts in the sensory systems (Vieira and Rozas 2011; Missbach et al. 2015; D. Wang et al. 2018; Almudi et al. 2020; Anholt 2020). Subterranean specialization has also offered opportunities for evolutionary innovation in the way animals interact with this particular environment (Cartwright et al. 2017). While adapting to the subterranean niches, different species, ranging from fishes to insects, have evolved highly convergent alternatives to live into perpetual darkness in habitats exhibiting specific biotic and abiotic factors (i.e., limited and heterogeneous nutrient sources, lack of primary production, constant temperature and humidity). Evolutionary regressions (e.g., loss of eyes and pigmentation), elaborated elements (e.g., hypertrophy of extra-optic sensory structures) and other physiological changes (e.g., loss of circadian rhythms and modified life cycles) have been reported as possible adaptations for many obligate subterranean fauna (Pipan and Culver 2012). Likewise, it is conceivable that the subterranean selective pressures have driven adaptive shifts in other sensory systems, including the chemosensory repertoires of subterranean animals. For instance, some studies on cavefish pointed out an enhancement of chemosensory systems from a morphological point of view (i.e., visible differences in taste buds and olfactory neural bulbs) when compared to surface populations (Parzefall 2001; Yamamoto et al. 2009; Yang et al. 2016). In subterranean arthropods, elongation of antennae and body appendages have been also attributed to enhanced sensory capabilities (Turk et al. 1996). Nevertheless, the evolution of the chemosensory repertoire in subterranean fauna from a molecular perspective remains widely unexplored.

Environmental chemical signals are enormously diverse in nature. Animals have developed a wide diversity of mechanisms to perceive and interpret specific cues essential to their evolutionary success (Nei et al. 2008). In insects, these chemicals comprehend palatable nutrient or repellent odors and tastes, pheromones, warning signals of predators and those indicating optimal substrates for oviposition, and various others (Joseph and Carlson 2015). The chemosensory system in insects is distributed morphologically in the interface between the environment and the dendrites of the peripheral sensory neurons, where different chemosensory proteins act in parallel for the signal transduction to the brain centers in which the information is processed (Joseph and Carlson 2015; Dippel et al. 2016). To capture this complex information, insects encode three large and divergent families of transmembrane chemoreceptor proteins: gustatory receptors (GRs), odorant receptors (ORs) and ionotropic receptors (IRs) (Clyne et al. 1999; Gao and Chess 1999; Vosshall et al. 1999; Benton et al. 2009; Sánchez-Gracia et al. 2009). GRs, which detect non-volatile compounds, likely represent the oldest chemosensory receptors (Eyun et al. 2017), being distributed in several taste organs along the entire body including mouth pieces, legs, wing margins and other specialized structures such as vaginal plate sensilla in female flies abdomen (Stocker 1994). Airborne chemical particles are perceived in the head appendages by the ORs, an insect-specific chemoreception gene family thought to have originated from the GR gene family (Robertson et al. 2003; Robertson 2019; Thoma et al. 2019). ORs work with the functionally essential and highly conserved odorant receptor co-receptor (ORCO), which was proposed as the ancestral OR. Moreover, IRs derived from the ionotropic glutamate receptor genes (IGluRs) superfamily in protostomes (Vosshall and Stocker 2007; Benton 2015) and mediate responses to many organic acids and amines, including pheromones and nutrient odors (Benton et al. 2009). In insects there are other gene families that also participate in the chemosensory function, such as the sensory neuron membrane proteins (SNMPs) (Grimaldi and Engel 2005; Nichols and Vogt 2008; Missbach et al. 2014). The odorant-binding proteins (OBPs) and chemosensory proteins (CSPs) also play a key role for chemoreception in terrestrial insects, besides other physiological roles. The stable and compact structure of OBPs and CSPs make them versatile soluble proteins relevant for signal transduction of small hydrophobic compounds such as pheromones and odorants (Roys 1954; Stürckow 1970; Pelosi et al. 2014; Pelosi et al. 2018).

Like in other large gene families encoding ecologically relevant proteins, constant birth-and-death dynamics may play an important role in their evolution in insects (Nei and Rooney 2005; Vieira et al. 2007; Sánchez-Gracia et al. 2009). A general positive correlation has been observed when comparing the chemosensory gene diversity across species and the chemical signals complexity of the ecological niche they occupy. Contrasting patterns of gene expansions and losses are found when exploring the chemosensory gene repertoires in extreme specialist and generalist species, with the latter usually exhibiting larger expansions of genes involved in chemoreception (Andersson et al. 2019; Robertson 2019). However, the evolution of the chemosensory gene families in subterranean species are still largely unexplored, hampering our understanding on how these animals perceive their particular environment.

Cave beetles represent ideal models to shed light on the genomic basis of chemoreception in subterranean environments. The Leptodirini tribe is a speciose lineage of scavenger beetles that represents one of the most impressive radiations of subterranean organisms. Several lineages within Leptodirini (estimated to have colonized subterranean habitats ca. 33 Mya; Ribera et al. 2010) acquired morphological and physiological traits typically associated with troglobitic adaptations. Their modifications include complete lack of eyes and optic lobes, depigmentation, membranous wings, elongation of antennae and legs (Jeannel 1924; Deleurance 1963; Luo et al. 2019) and loss of thermal acclimation capacity (Rizzo et al. 2015; Pallarés et al. 2018). They also exhibit singular modified life cycles as a key innovation for their subterranean specialization (Delay 1978; Cieslak et al. 2014). One of the highly modified species of the Leptodirini tribe is *Speonomus longicornis* Saulcy, 1872 (Coleoptera, Polyphaga, Leiodidae) (fig. 1a). This obligate cave-dwelling beetle is completely blind, depigmented, possesses enlarged antennae (Jeannel 1924) with a high sensilla density (fig. 1b) and it has a contracted life cycle, comprising a single larval-instar during its development in which the larvae remains practically quiescent like the pupal stage (Glaçon 1953). The troglobitic characters of this species suggest an extreme adaptation to the deep subterranean.

**Figure 1.**
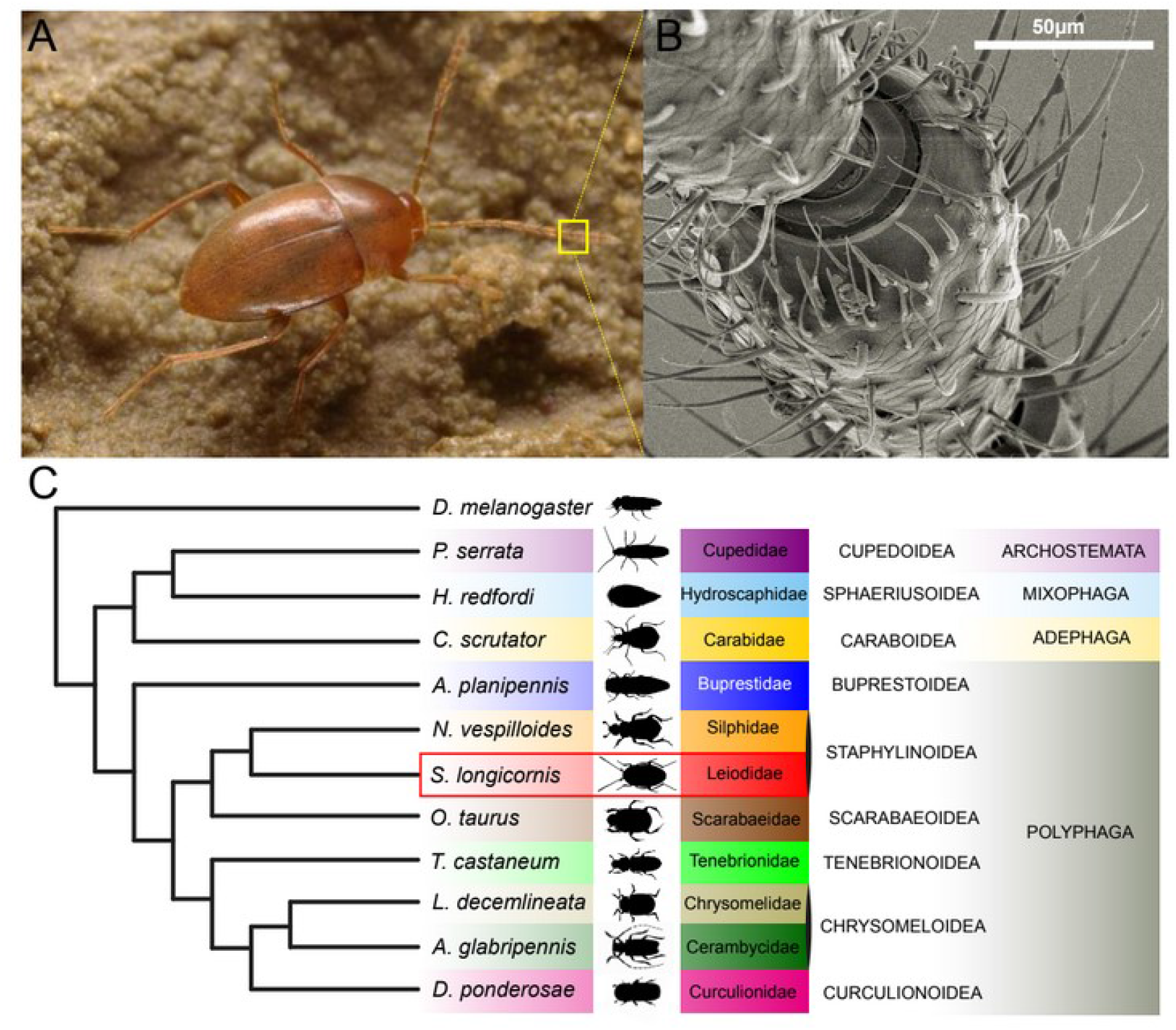
(A) *Speonomus longicornis.* (B) Scanning Electron Microscope image of the antennal sensilla of *S. longicornis* (voucher IBE-AI531). (C) Simplified phylogeny showing the relationships of the studied species, adapted from (McKenna et al. 2019)).

The present study aims to characterize the chemosensory gene repertoire of *S. longicornis*. The goals of this project are (i) to pinpoint genes putatively involved in chemoreception in the cave beetle *S. longicornis* through a transcriptomic approach, and (ii) to explore how such genes evolved in the broader phylogenetic context of beetle and insect evolution. Our study therefore aims at providing the first characterization of the chemosensory gene repertoire of an obligate cave-dwelling species.

## MATERIALS AND METHODS

### Sample collection and preservation

Thirty specimens of *Speonomus longicornis* were collected in 2016 at the type locality: Grotte de Portel cave, in the Plantaurel massif at the French region of Ariège (43°01’51”N, 1°32’22’E). All specimens were manually captured and kept alive inside a thermo-box during the stay at the cave. Once the sampling was finished, all individuals were placed in an 8 ml tube and flash-frozen in liquid nitrogen at the cave entrance in order to prevent stress-related alterations in the expression levels and to minimize RNA degradation during transportation to the laboratory, where the samples were stored at −80°C until the RNA extraction.

### RNA extraction

All steps were performed in cold and RNase free conditions. Several specimens were pooled in each sample in order to obtain sufficient tissue for an efficient extraction. We did not examine the sex of the specimens to minimize the manipulation in order to avoid RNA degradation. Nevertheless, no significant sexual dimorphism has been found in the chemosensory system of other coleopterans (Dippel et al. 2016).

The specimens were split in three groups of ten individuals each, representing biological replicates. Since chemosensory structures are mainly concentrated in the antennae (see Introduction), we dissected the antennae of each specimen. Therefore, our experimental design included 3 biological replicates representing 2 conditions: antennae and the rest of the body.

The isolation of total RNA was performed by Phenol/Chloroform extraction, with a lysis through guanidinium thiocyanate buffer following the protocol of Sambrook et al. (1989) with minor modifications (i.e., not using 2-mercaptoethanol). A first quality check was done by size separation in a 1% TBE agarose gel chromatography. Total RNA yield was quantified by a RNA assay in a Qubit fluorometer (Life Technologies).

### cDNA Library Construction and Next-Generation Sequencing

For the antennae samples, a low-input RNA sequencing protocol was followed. mRNA sequencing libraries were prepared following the SMARTseq2 protocol (Picelli et al. 2013) with some modifications. Briefly, RNA was quantified using the Qubit® RNA HS Assay Kit (Thermo Fisher Scientific). Reverse transcription with the input material of 2ng was performed using SuperScrpit II (Invitrogen) in the presence of oligo-dT30VN (1μM; 5-AAGCAGTGGTATCAACGCAGAGTACT30VN-3’), template-switching oligonucleotides (1μM) and betaine (1M). The cDNA was amplified using the KAPA Hifi Hotstart ReadyMix (Roche), 100 nM ISPCR primer (5’-AAGCAGTGGTATCAACGCAGAGT-3’) and 15 cycles of amplification. Following purification with Agencourt Ampure XP beads (1:1 ratio; Beckmann Coulter), product size distribution and quantity were assessed on a Bioanalyzer High Sensitivity DNA Kit (Agilent). The amplified cDNA (200 ng) was fragmented for 10 min at 55 °C using Nextera® XT (Illumina) and amplified for 12 cycles with indexed Nextera® PCR primers. The library was purified twice with Agencourt Ampure XP beads (0.8:1 ratio) and quantified on a Bioanalyzer using a High Sensitivity DNA Kit.

For the samples containing the rest of the body, total RNA was assayed for quantity and quality using the Qubit® RNA BR Assay kit (Thermo Fisher Scientific) and RNA 6000 Nano Assay on a Bioanalyzer 2100 (Agilent). The RNASeq libraries were prepared from total RNA using the KAPA Stranded mRNA-Seq Kit for Illumina (Roche) with minor modifications. Briefly, after poly-A based mRNA enrichment from 500ng of total RNA, the mRNA was fragmented. The second strand cDNA synthesis was performed in the presence of dUTP instead of dTTP, to achieve strand specificity. The blunt-ended double stranded cDNA was 3’adenylated and Illumina single indexed adapters (Illumina) were ligated. The ligation product was enriched with 15 PCR cycles and the final library was validated on an Agilent 2100 Bioanalyzer with the DNA 7500 assay.

The libraries were sequenced on an Illumina HiSeq 2500 platform in paired-end mode with a read length of 2×76bp. Image analysis, base calling and quality scoring of the run were processed using the manufacturer’s software Real Time Analysis (RTA 1.18.66.3) and followed by generation of FASTQ sequence files by CASAVA. cDNA libraries and mRNA sequencing were performed at the National Center of Genomic Analyses (CNAG) (Barcelona, Spain).

### Sequence Processing, decontamination and *de Novo* Assembly

Raw reads for all samples were downloaded in FASTQ format. The quality of the raw reads was assessed and visualized using FASTQC v.0.11.8 (www.bioinformatics.babraham.ac.uk). For each dataset, remaining Illumina adaptors were removed and low-quality bases were trimmed off according to a threshold average quality score of 30 based on a Phred scale with Trimmomatic v. 0.38 (Bolger et al. 2014). Filtered paired-end reads were validated through a FASTQC visualization.

A reference *de novo* transcriptome assembly was constructed with Trinity v.2.8.4, using paired read files and default parameters, including all replicates and conditions (Grabherr et al. 2011; Haas et al. 2013). Blobtools v.1.1.1 (Laetsch and Blaxter 2017) was used to detect putative contamination from the assembled transcriptome. Transcripts were annotated using BLAST+ v.2.4.0 against the non-redundant (nr) database from NCBI with an expected value (E-value) cutoff of 1e^-10^; and mapped to the reference transcriptome with Bowtie2 v.2.3.5.1 (Langmead and Salzberg 2012). Putative contaminants included transcripts with significant hits to viruses, fungi, bacteria or chordates, accounting for a total of 7.8% of the mapped sequences (see also supplementary figure S1).

### Inference of candidate coding regions and transcriptome completeness assessment

To check completeness of the reference transcriptome, we searched for single copy universal genes in arthropods. For that, we used BUSCO v.2.0.1 (Benchmarking Universal Single-Copy Orthologs; (Simão et al. 2015) using the arthropoda database (arthropoda_odb9) through the gVolante web server with default settings (Nishimura et al. 2017). The assembly was processed in TransDecoder v.5.4.0 to identify candidate open reading frames (ORFs) within the transcripts using the universal genetic code and a minimum length of 100 amino acids (Haas et al. 2013). Only the longest ORFs were retained as final candidate coding regions for further analyses.

### Chemosensory gene repertoire characterization

Bitacora v.1.0.0 (Vizueta et al. 2020) was used to curate annotations during the sequence similarity searches of the chemosensory gene families of interest. Curated protein databases containing chemoreceptor genes (ORs, GRs, IRs, SNMPs, OBPs and CSPs) of several arthropods were used in the Bitacora searches (Vizueta et al. 2016). For the ORs annotation, we also used an additional database containing Coleoptera ORs based on the datasets from Mitchell et al. (2019). The “protein mode” pipeline of Bitacora was used to annotate the ORFs of the transcriptome, combining BLAST and HMMER searches. All the predicted coding regions were implemented for Bitacora searches, retrieving a multifasta file for each of the chemosensory families. Results were filtered with customized Python scripts using Biopython v.1.76 SeqIO package (https://biopython.org; (Cock et al. 2009); in order to reduce redundancy and to validate dubious annotations (i.e., some ORFs received significant hits for both ORs and GRs, which were post-validated through Pfam searches). To reduce redundancy, final candidates were represented by the longest isoform per gene and thus achieving unique gene annotations.

### Expression levels quantification and differential gene expression analysis

Salmon v. 0.10.2 (Patro et al. 2017) was used for indexing and quantification of transcript expression. Expression estimated counts were transformed into an expression matrix using a Perl script included in the Trinity software (abundance_estimates_to_matrix.pl), which implements the trimmed mean of M-values normalization method (TMM). We also explored the variability of the estimated expression values within and between replicates and samples using several Perl scripts included in Trinity v. 2.8.4 analysis toolkit. Differential expression analysis was conducted in the Bioconductor edgeR package (Robinson et al. 2010; Robinson and Oshlack 2010). The Benjamini-Hochberg method was applied to control the false discovery rate (FDR) (Benjamini and Hochberg 1995). Significance value for multiple comparisons was adjusted to 0.001 FDR threshold cutoff and a 4-fold change. Differentially expressed genes (upregulated and downregulated) in antennae and the rest of the body were plotted in scatter plots, MA plots and heatmaps using the R scripts provided in the Trinity software. The expression matrix was interrogated in order to detect exclusively expressed genes in antennae (defined as genes that showed positive expression values in the three replicates of antennae and with expression values lower than 0.001 in the rest of the body).

### Transcriptome characterization, Gene Ontology (GO) enrichment and visualization

The peptide predictions, including all isoforms, were used as input for eggNOG-mapper v.4.5.1 (Huerta-Cepas et al. 2017), retrieving Gene Ontology (GO) terms for all the annotated transcripts. The GO annotations were subsequently filtered to discard those corresponding to non-animal taxa (i.e., viruses, bacteria, fungi, plants, 10.86% of total GO annotations) and to eliminate the redundancy provided by the isoforms. All GO terms for each unique gene were retained. GO enrichment analysis was performed using the Fisher’s test in FatiGO software (Al-Shahrour et al. 2007) to detect significant over-representation of the GO terms in the pairwise comparisons between the upregulated genes in antennae and the rest of the body, adjusting the p-value to 0.05.

GO enrichment results were visualized in the REVIGO web server (Supek et al. 2011), plotting the results in a “TREEMAP” graph using R, where the size of the rectangles is proportional to the enrichment p-value (abs_log10_pvalue) of the overrepresented GO terms.

### Phylogenetic inferences for the candidate chemosensory genes

Complete and partial annotated genes for *S. longicornis* (referred to as *Slon* in the figures) were included in the phylogenetic inferences in order to interrogate their phylogenetic relationships with chemosensory genes of other species, all of them based on genomic data. With this approach, we aim to infer diversity patterns of the chemosensory repertoire of *S. longicornis* and to characterize each gene family more specifically, indicating with a higher confidence the putative function of these genes compared to analysis merely based on homology. Since no reference genome is available for the focus species, we inferred a deeply-sequenced de novo transcriptome which resulted in a high completeness based on our assessment of BUSCO genes (see Results), indicating that we recovered a mostly complete reference gene set and hence it is of enough quality to explore gene family evolution (Cheon et al. 2020). This approach has been successfully applied in other studies with non-model organisms through the combination of genomic and high quality transcriptomic data (e.g., Fernández and Gabaldón 2020; Vizueta et al. 2020).

Individual phylogenies for each chemosensory gene family as annotated by Bitacora (see above) were inferred using the following pipeline. Amino acid sequences were aligned using PASTA software v.1.7.8 (Mirarab et al. 2015). Poorly aligned regions were trimmed using trimAl v.1.2 (Capella-Gutiérrez et al. 2009) with the ‘-automated1’ flag. Maximum likelihood phylogenetic inference was inferred with IQ-TREE v.2.0.4 (Nguyen et al. 2015). We applied the mixture model LG+C20+F+G with the site-specific posterior mean frequency model (PMSF; (H.-C. Wang et al. 2018) and the ultrafast bootstrap option (Hoang et al. 2018). A guide tree was inferred with FastTree2 under the LG model (Price et al. 2010). Results were visualized using the iTOL web interface (Letunic and Bork 2019).

The ORs phylogeny included the coleopteran ORs obtained from (Mitchell et al. 2019) (fig. 1c). These species included a range of ecological strategies, as follows. The phytophagous specialists were represented by the aquatic beetle *Hydroscapha redfordi* (Myxophaga, Hydroscaphidae), the ash borer *Agrilus planipennis* (Polyphaga, Buprestidae), the Colorado potato beetle *Leptinotarsa decemlineata* (Polyphaga, Chrysomelidae) and the mountain pine beetle *Dendroctonus ponderosae* (Polyphaga, Curculionidae). Moreover, the species set include two non-phytophagous beetles, the insectivorous *Calosoma scrutator* (Adephaga, Carabidae) and the burying beetle *Nicrophorus vespilloides* (Polyphaga, Silphidae), which feeds on carrion. Hence, both these phytophagous and non-phytophagous species can be considered as oligophagous, although the phytophagous species are strict specialists and the non-phytophagous are not. By contrast, the generalist species included the dung beetle *Onthophagus taurus* (Polyphaga, Scarabaeidae), the red flour beetle *Tribolium castaneum* (Polyphaga, Tenebrionidae), the wood borer *Anoplophora glabripennis* (Polyphaga, Cerambycidae) and the reticulated beetle *Priacma serrata* (Archostemata, Cupedidae), that is presumed to feed on seeds (Hörnschemeyer et al. 2013). For the ORCOs phylogeny, we also included some additional ORCO sequences of coleopteran species and other taxa as outgroups *(Apis mellifera* and *D. melanogaster)* (see species and GeneBank accessions at supplementary table S1).

For the remaining gene families explored in this study (GRs, IRs, IGluRs, SNMPs, OBPs and CSPs), we selected a custom taxon set for each case based on sequence availability of high quality annotated genes in previous studies. For most gene families, we included sequences of *T. castaneum, D. ponderosae, A. planipennis.* Several functionally characterized GRs, IRs, IGluRs and SNMPs of *Drosophila melanogaster* obtained from FlyBase (https://flybase.org), were also included in order to characterize those genes in *S. longicornis. T. castaneum* sequences were retrieved from Dippel et al., (2014) (OBPs and CSPs) and Dippel et al., (2016) (GRs, IRs and SNMPs) and translated to amino acid sequences prior to alignment, trimming and phylogenetic analyses, following the same pipeline as described above. IRs, GRs, SNMPs, OBPs and CSPs of *D. ponderosae* and *A. planipennis* were acquired from Andersson et al (2019). IR sequences from Croset et al. (2010) and Wang et al. (2015) representing several invertebrate phyla were retrieved to further explore phylogeny of the early splitting clades IR8a and IR25a at a larger evolutionary scale (suppl. table 1). This phylogeny includes genes of *D. melanogaster, Aedes aegypti, Culex quinquefasciatus, Anopheles gambiae, Bombyx mori, Apis mellifera, Nasonia vitripennis,* Microplitis mediator, *Acyrthosiphon pisum, Pediculus humanus, Daphnia pulex, Caenorhabditis elegans*, *Capitella capitata*, *Aplysia californica* and *Lottia gigantea*.

## RESULTS

### A high quality *de novo* transcriptome for *Speonomus longicornis* facilitates the annotation of its chemosensory gene repertoire

A *de novo* assembly transcriptome was constructed combining the pair-end reads from the six libraries (~411 millions of reads; ~356 millions after trimming), obtaining a total of 245,131 transcripts (supplementary figure S1). These transcripts include 177,711 unique predicted ‘genes’ by Trinity and 74,273 candidate open reading frames (ORFs, including all genes and isoforms). When filtering by the longest isoform per gene (which could be considered as a ‘proxy’ for the total number of genes that we may encounter in the genome), we obtained a total of 20,956 ORFs. BUSCO analysis indicated a high completeness for the assembled transcriptome, with 99% of complete BUSCO genes compared to the Arthropoda database. Further details about sequencing, assembly statistics, the completeness assessment and the putative contamination results are summarized in the supplementary fig. S1 (see Supplementary Material online).

### Differential gene expression analysis reveals chemosensory genes upregulated in the antennae

Bitacora searches identified a total of 205 chemosensory gene candidates for *S. longicornis* (Table 1, see also supplementary table S2). The expression level distribution obtained in the transcript quantification steps was supervised in order to identify possible biases when comparing replicates and conditions (supplementary figure S2). Our results indicate that replicates are more similar to each other than between the different conditions. We detected 18,160 clusters of transcripts (reported as clusters of transcripts or ‘genes’ by Trinity, referred to as Trinity genes hereafter) differentially expressed in antennae and body, with 8,949 Trinity genes upregulated in antennae (supplementary figure S3 and supplementary table S3). Out of the 205 candidate chemosensory genes as detected by Bitacora, 78 were detected as differentially expressed. From those, 49 genes were overexpressed in antennae (18 ORs, 17 OBPs, 5 GRs, 5 IRs/IGluRs, 2 SNMPs/CD36s and 2 CSPs) (fig. 2A and table 1). We also identified 7 ORs exclusively expressed in antennae, including 2 additional genes not recovered as differentially expressed due to the disparity of the expression values between antennae replicates (supplementary table S4).

**Figure 2.**
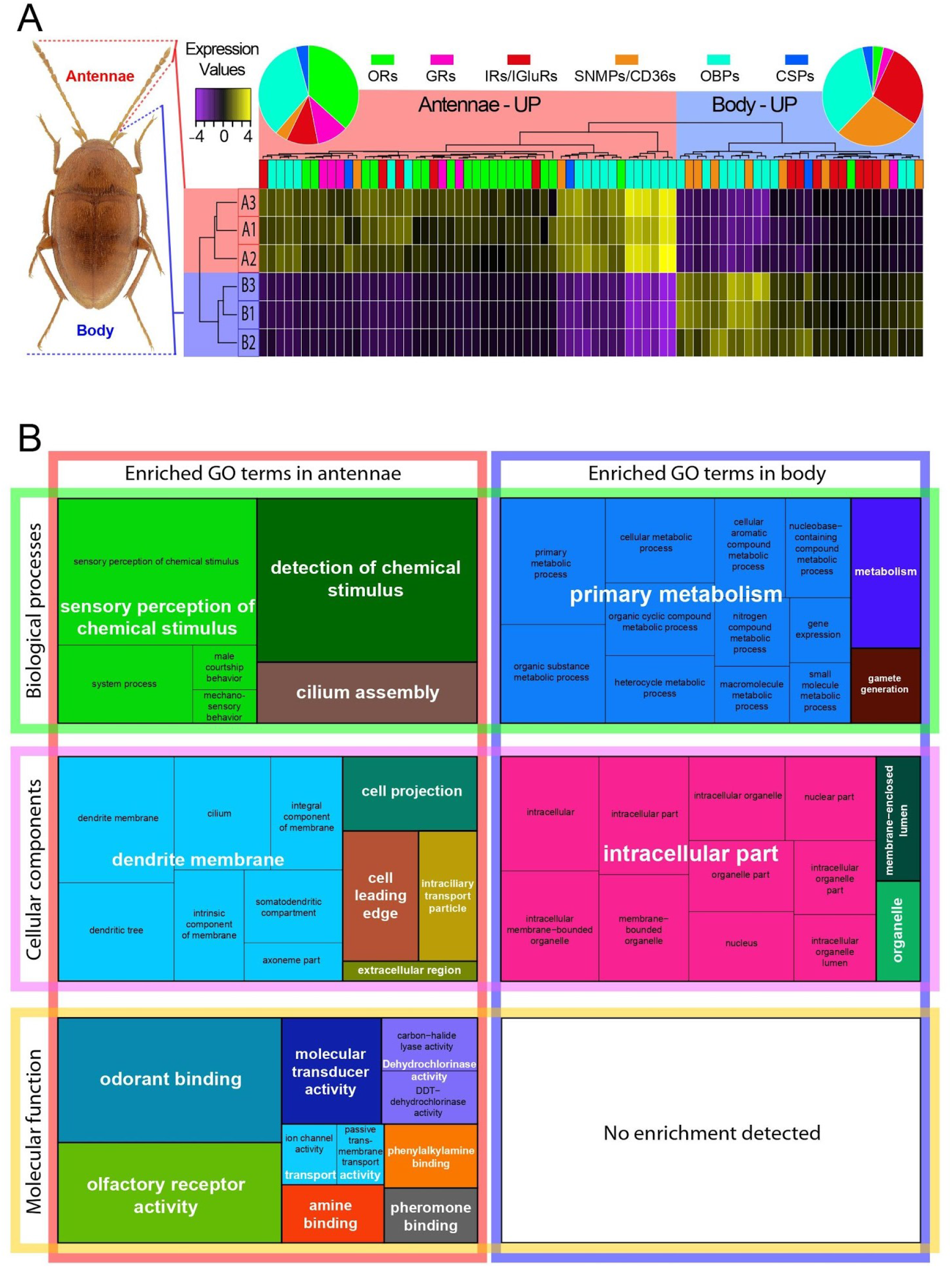
(A) Heatmap of chemosensory genes of *S. longicornis* differentially expressed in antennae and the rest of the body. (B) Gene ontology (GO) treemaps for the differentially expressed genes in antennae versus the rest of the body. Biological process, molecular function and cellular component enriched GO terms are shown.

**Table 1.**
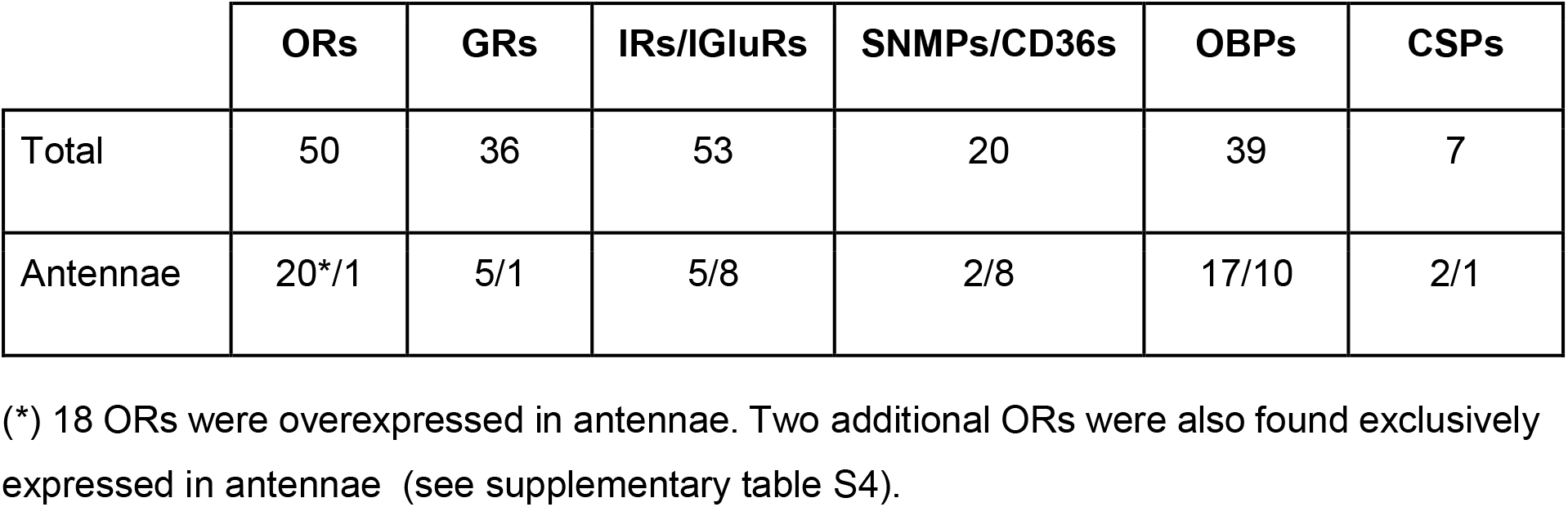
Number of annotated chemosensory genes of *S. longicornis* and number of overexpressed / underexpressed genes in antennae.

### Gene Ontology enrichment reveals upregulated chemosensory specificity in antennae

Out of the 44,107 annotated ORFs, only 23,555 had associated GO terms, representing 31.7% of the total queried sequences from the assembled transcriptome. After filtering annotations from non-animal taxa (including viruses, bacteria and fungi), we retained 89.14% of the annotations. Figure 2B depicts enriched GO terms for upregulated genes in antennae and in the rest of the body. In antennae, “sensation and perception of chemical stimulus” represent the most enriched category within the biological processes analysed and, in less proportion, some categories related to cilium activity. “Mechanosensory activity” terms are also overrepresented but in a minor proportion. More than half of the cellular components GOs enriched in antennae correspond to “dendritic structures”, and a high proportion of the overrepresented terms correspond to “extracellular and membrane structures”. Regarding the molecular function category in antennae, “odorant-binding” and “odorant reception” terms occupy a large proportion of the enriched functions followed by other “binding and signal transduction” terms.

### Phylogenetic interrelationships of Coleoptera ORs and ORCOs

A total of 1,222 OR sequences were aligned and trimmed (see Material and Methods). The final length of the alignment was of 254 amino acid positions. To facilitate comparison, we retained the nomenclature used by Mitchell et al. (2019) to describe the phylogenetic groups and clades recovered in their phylogenetic analyses (i.e., groups 1, 2A, 2B, 3, 4, 5A, 5B, 6 and 7). Our results were overall congruent with those reported by Mitchell et al. (2019), with virtually all OR groups recovered with high support except group 6 and a different position for group 4, which was recovered as nested within group 3 (fig. 3A). While most of the genes fall into the same groups than in Mitchell et al. (2019), four genes (i.e., AplaOR1, PserOR120-121 and SlonOR34525c0g1) were not recovered for any of the previously proposed groups. The upregulated ORs in *S. longicornis* antennae were distributed along the different coleopteran OR groups, mostly clustered within group 1 and group 7 (with 6 and 5 genes respectively). Exclusively expressed ORs in antennae are found in groups 1, 3, 4 and 7. The number of ORs is highly variable among these species (fig. 3B). *S. longicornis* and the other non-phytophagous species (i.e*. C. scutator, N. vespilloides)* exhibited relatively moderate OR repertoires and a similar distribution pattern (i.e., without representation in group 5A, moderate gene expansions and finding their largest expansion in group 3; fig 3B and 3D). All the ORCO sequences were recovered as a clade with high support and were used to root the tree, facilitating the identification of the ORCO candidate of *S. longicornis* (SlonORCO; fig. 3A and 3C).

**Figure 3.**
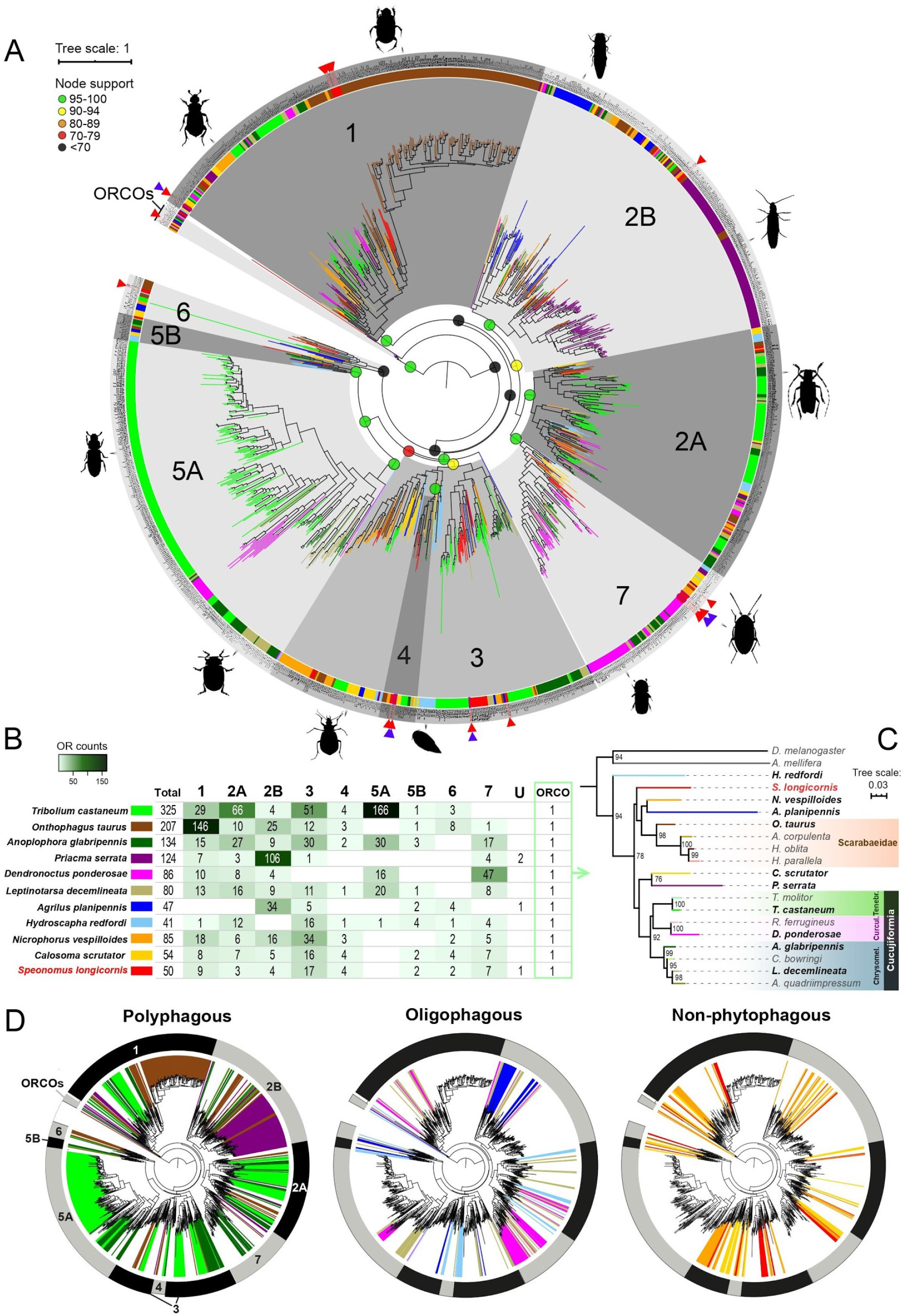
(A) Maximum likelihood phylogenetic tree of odorant receptors (ORs) including OR sets of *S. longicornis* and other coleopterans from Mitchell et al. (2019), representing the proposed OR groups in grey ranges. Red triangles represent the upregulated genes in the antennae of *S. longicornis*. Purple triangles represent exclusively expressed genes in antennae. Species are color coded as indicated in fig. 3B. (B) Number of OR genes of each OR group inferred for each species included in the phylogeny. ‘U’ indicates unclassified ORs. (C) Maximum likelihood phylogeny of ORCO across coleopterans (see methods and supplementary table S1 for species codes). (D) Simplified representation of OR diversity recovered for each species, highlighting the OR repertoire of species with different feeding strategies.

The phylogeny of ORCOs (fig. 3C) (with a final trimmed alignment of 478 amino acid positions) recovered clades for the different beetle families, but did not mirror the phylogeny of Coleoptera at the family level. For instance, all ORCOs of species of Cucujiformia were recovered in a clade and were subsequently clustered into their corresponding families. The same pattern was observed for the ORCOs of the different species of Scarabaeidae. By contrast, the ORCO of *S. longicornis* did not cluster together with that of *N. vespilloides*, despite belonging to the same superfamily (i.e., Staphylinoidea).

### Phylogenetic inference of the annotated GRs

A total of 374 sequences were interrogated, resulting in a multiple sequence alignment of 271 amino acid positions. Notably, none of the candidate GRs from *S. longicornis* clustered together with GRs involved in the reception of fructose and other sugars in the other species (fig. 4). On the other hand, in several coleopteran GRs, including *S. longicornis,* we recovered 3 candidates that cluster with those of *D. melanogaster* involved in perception of CO_2_, which are well characterized functionally (termed as GR1, GR2, GR3 in beetles, and GR21a, GR63a in *D. melanogaster*; (Jones et al. 2007; Kwon et al. 2007; Dippel et al. 2016). One of these three candidate CO_2_ receptors of *S. longicornis* was upregulated in antennae, that was recovered as orthologous to the GR2 gene in beetles. We also identified a candidate bitter taste GR (SlonGR19567c0g1) that clustered together with strong support with previously identified as conserved bitter taste GRs for *A. planipennis* and *D. ponderosae* (Andersson et al. 2019). The rest of the genes were generally recovered in well-supported clades with species-specific differences in the extent of GR expansions. *S. longicornis* showed divergent GRs distributed along the tree exhibiting relatively small expansions (i.e., from two to five genes) and in general a relatively diminished gustatory repertoire, whereas the other species exhibit remarkable expansions and considerably larger repertoires. The functions of the other four upregulated GRs in antennae cannot be further characterized due to the lack of functional annotation of the genes they cluster together with.

**Figure 4.**
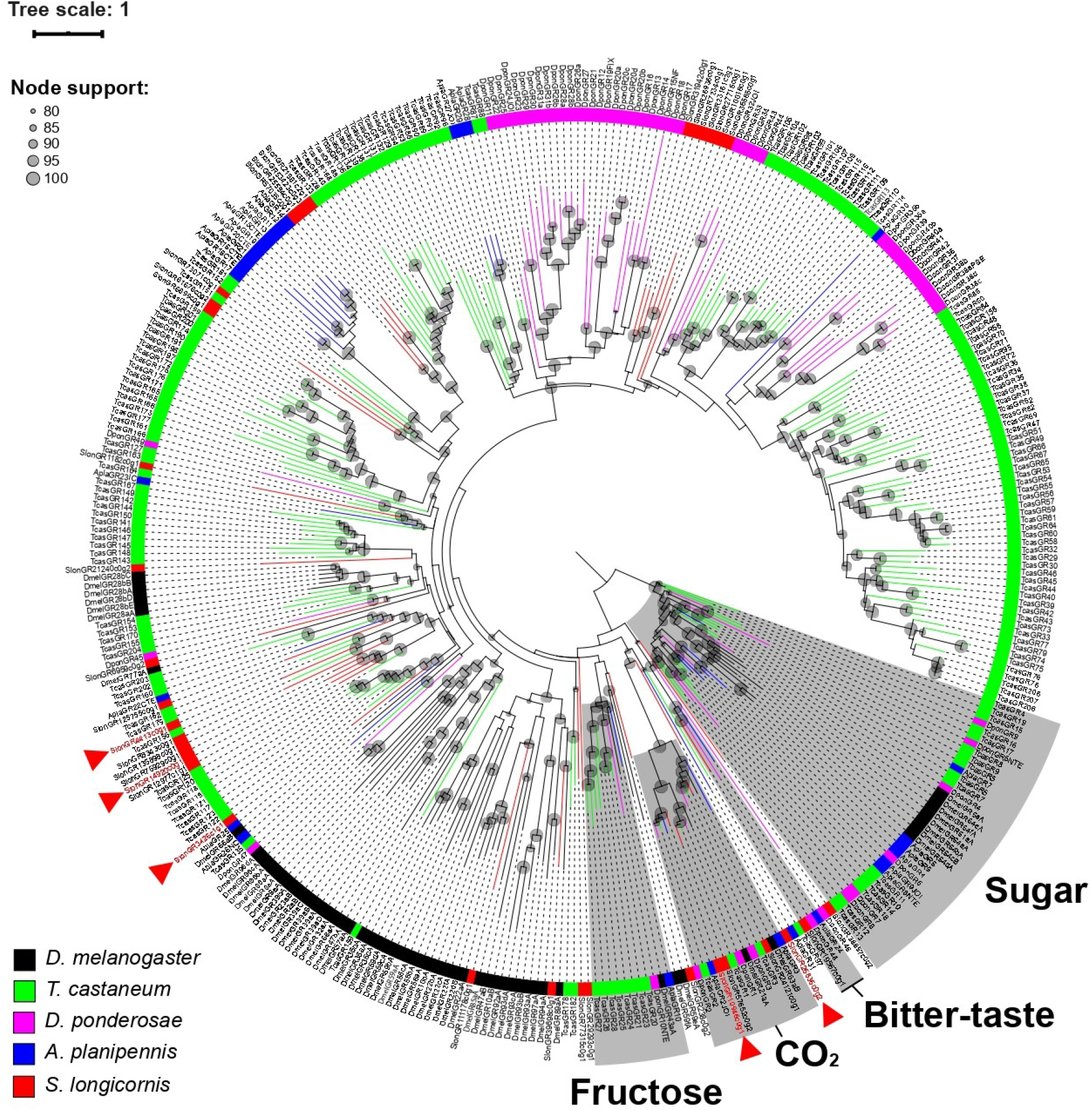
Maximum likelihood phylogenetic tree of gustatory receptors (GRs) including GR sets of *S. longicornis*, other coleopterans from Andersson et al. (2019) and conserved GR sequences of *D. melanogaster*. Grey ranges represent well supported GR clades, indicating the proposed functions in the other species. Red triangles represent upregulated genes in the antennae of *S. longicornis*.

### IRs and IGluRs phylogenies

Since IRs derive from IGluRs (see Introduction), our phylogenetic approach facilitated the initial annotations of both types of genes in *S. longicornis.* IGluRs are not directly associated with chemoreception but present a high sequence identity with the most conserved IRs (i.e., IR8a, IR25a). Three genes clustered together with N-methyl-D-aspartate receptors (NMDARs) and nine genes clustered with different IGluR clades, one upregulated in antennae (fig. 5A).

**Figure 5.**
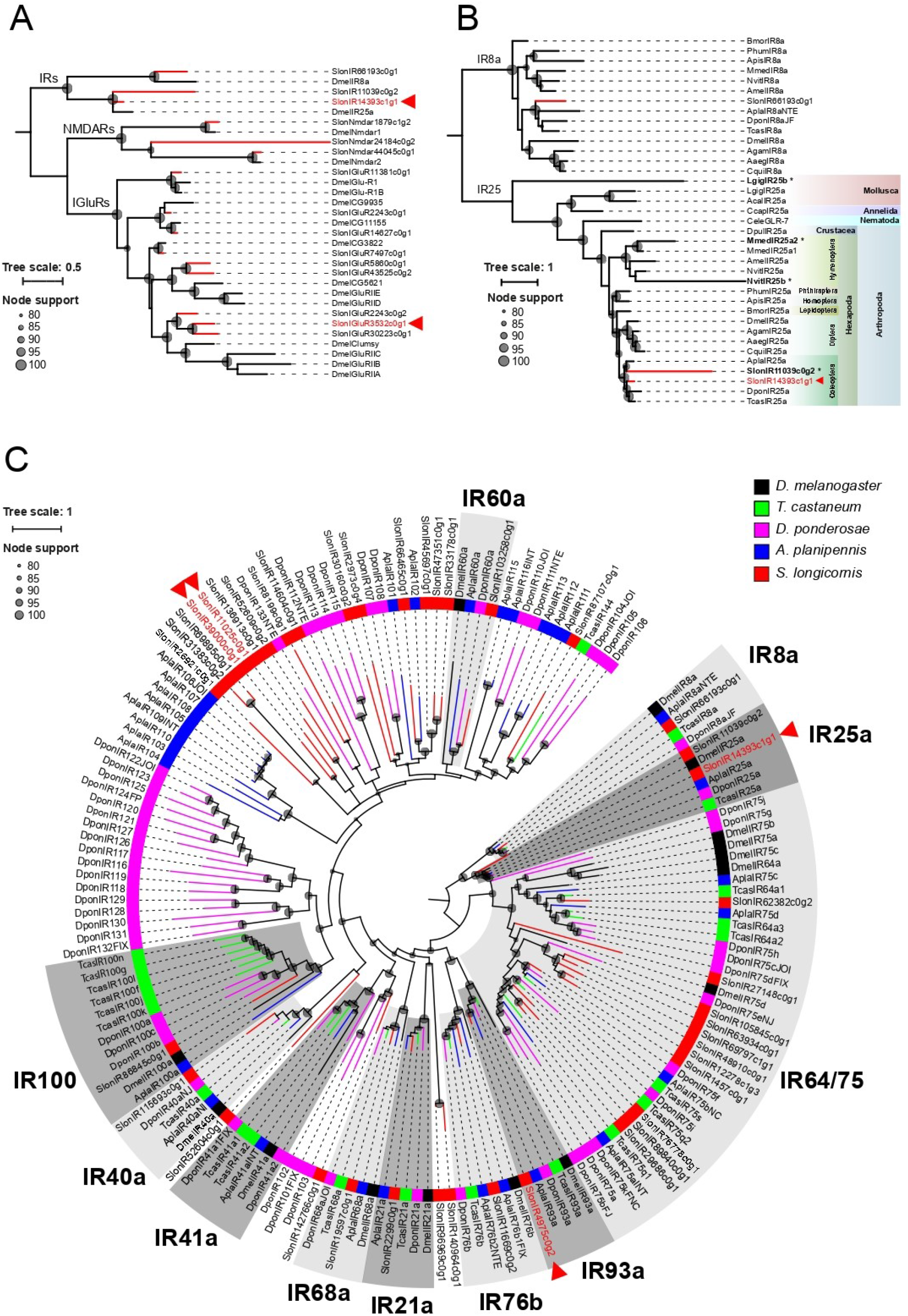
(A) Maximum likelihood phylogenetic tree of ionotropic glutamate receptors (IGluRs) including sequences of *S. longicornis* and *D. melanogaster*. (B) Maximum likelihood phylogenetic tree of the ionotropic receptors clades IR25a and IR8a (the later used to root the tree), including the candidate genes for *S. longicornis* (*Slon*), beetle sequences retrieved from Andersson et al. (2019) and Dippel et al. (2016) and sequences from an expanded invertebrate taxon sampling obtained from Croset et al. (2010) and Wang et al. (2015). Species codes are as follows: *D. melanogaster* (*Dmel*), *Aedes aegypti* (*Aaeg*), *Culex quinquefasciatus* (*Cqui*), *Anopheles gambiae* (*Agam*), *Bombyx mori* (*Bmor*), *Apis mellifera* (*Amel*), *Nasonia vitripennis* (*Nvit*), Microplitis mediator *(Mmed) Acyrthosiphon pisum* (*Apis*), *Pediculus humanus* (*Phum*), *Daphnia pulex* (*Dpul*), *Caenorhabditis elegans* (*Cele*), *Capitella capitata* (*Ccap*), *Aplysia californica* (*Acal*) *and Lottia gigantea* (*Lgig*). Asterisks indicate the divergent copies of IR25 candidates. (C) Maximum likelihood phylogenetic tree of ionotropic receptors (IRs) including IR sequences of *S. longicornis,* other coleopterans from Andersson et al. (2019) and IR sequences of *D. melanogaster.* Grey ranges represent the conserved IR clades, based on the annotations of the other species. In all trees, red triangles represent upregulated genes in the antennae of *S. longicornis*.

A total of 164 IR sequences were aligned and trimmed resulting in a multiple sequence alignment of 356 amino acid positions (fig. 5C). Several IRs of *S. longicornis* clustered together with conserved IRs in insects (i.e., IR8a, IR25a, IR93a, IR76b, IR21a, IR68a, IR40a, IR100, IR60a). No genes clustering together with IR41a were detected for *S. longicornis.* Several putative gene duplications were detected for *S. longicornis* (containing each from 2 to 5 IRs), more moderate in size than the large expansions observed for *A. planipennis* and *D. ponderosae,* which included up to 8 and 17 IRs, respectively. Our results detected a gene duplication in two genes clustering together with IR25a, a highly conserved single copy gene virtually in all protostomes (with the exception of the parasitoid wasps *N. vitripennis* and *M. mediator* and the limpet *L. gigantea,* see Discussion). The most conserved copy of IR25a in *S. longicornis* was upregulated in antennae (fig. 5A, 5B and 5C). In order to assess the robustness of our results, all isoforms from both genes were visually inspected in an alignment (supplementary figure S4), and the final alignment for the phylogeny including all the taxa (using the longest isoforms as described in Material and Methods) was examined to discard that they were nonoverlapping fragmented genes, confirming that this result may not be an artifact of our methodology. These IR25a candidates (i.e., SlonIR11039c0g2 and SlonIR14393c1g1) share 44% of identical residues whereas the conserved copy of *S. longicornis* (SlonIR14393c1g1) has between 69 to 72% of amino acid sequence identity with the IR25 candidates of the other coleopteran species. In addition, the protein annotation of the IR25a candidates of *S. longicornis* by HMMER resulted in highly similar domain profiles, suggesting their similarity at the structural level.

### SNMPs/CD36s phylogeny

A total of 41 sequences were aligned and trimmed resulting in a multiple sequence alignment of 382 amino acid positions (fig. 6). We followed the nomenclature used by Nichols and Vogt (2008) for ease of comparison (e.g., groups 1, 2 and 3). Genes involved in chemoreception were clearly identified as those belonging to group 3 of the SNMPs/CD36s clade (termed as SNMP1, 2 and 3 in insects), the only subfamily known to be involved in chemoreception in insects (Nichols and Vogt 2008). Our results indicated that S. *longicornis* SNMP candidate genes were formed by one SNMP1 gene, seven SNMP2 and one SNMP3, with a gene from SNMP1 upregulated in antennae. Remarkably, genes belonging to the subfamily SNMP2 were noticeably expanded compared to the other species, where they present only 1 or 2 genes (such as in *D. ponderosae*). Other candidate genes in *S. longicornis* within this gene family clustered together in groups 1 and 2, and include genes functionally annotated as scavenger receptors in *D. melanogaster* such as Croquemort (Crq), Peste (Pes), Santa Maria and ninaD (Nichols and Vogt 2008), with one gene from *S. longicornis* clustering with group 2 receptors and being upregulated in antennae.

**Figure 6.**
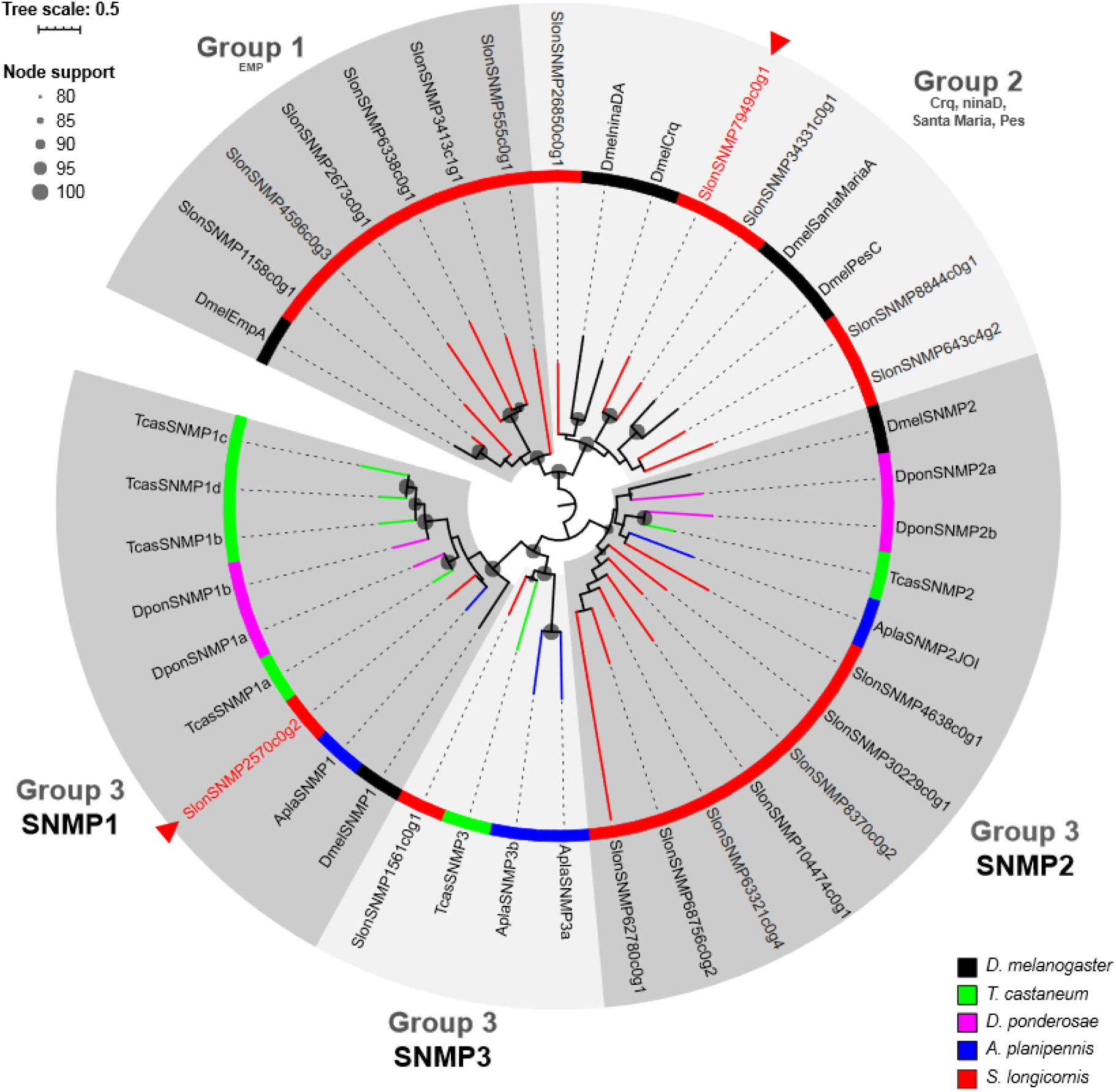
Maximum likelihood phylogenetic tree of sensory neuron membrane proteins (SNMPs/CD36s) including SNMP sequences of *S. longicornis,* other coleopterans from Andersson et al. (2019) and sequences of *D. melanogaster*. Grey ranges represent well supported SNMP/CD36 clades, which genes have been annotated in the other species. Red triangles represent upregulated genes in the antennae of *S. longicornis*.

### CSPs and OBPs phylogenies

For the CSPs phylogeny, we included a total of 52 sequences resulting in a multiple sequence alignment of 100 amino acid positions (fig. 7A). Unlike the CSP repertoires of the other species, *S. longicornis* does not present any duplication and it is relatively reduced. Moreover, we identified a candidate CSP clustered in the previously described as highly conserved CSPs clade in Coleoptera (DponCSP12, AplaCSP8, TcasCSP7E; Andersson et al. 2019).

**Figure 7.**
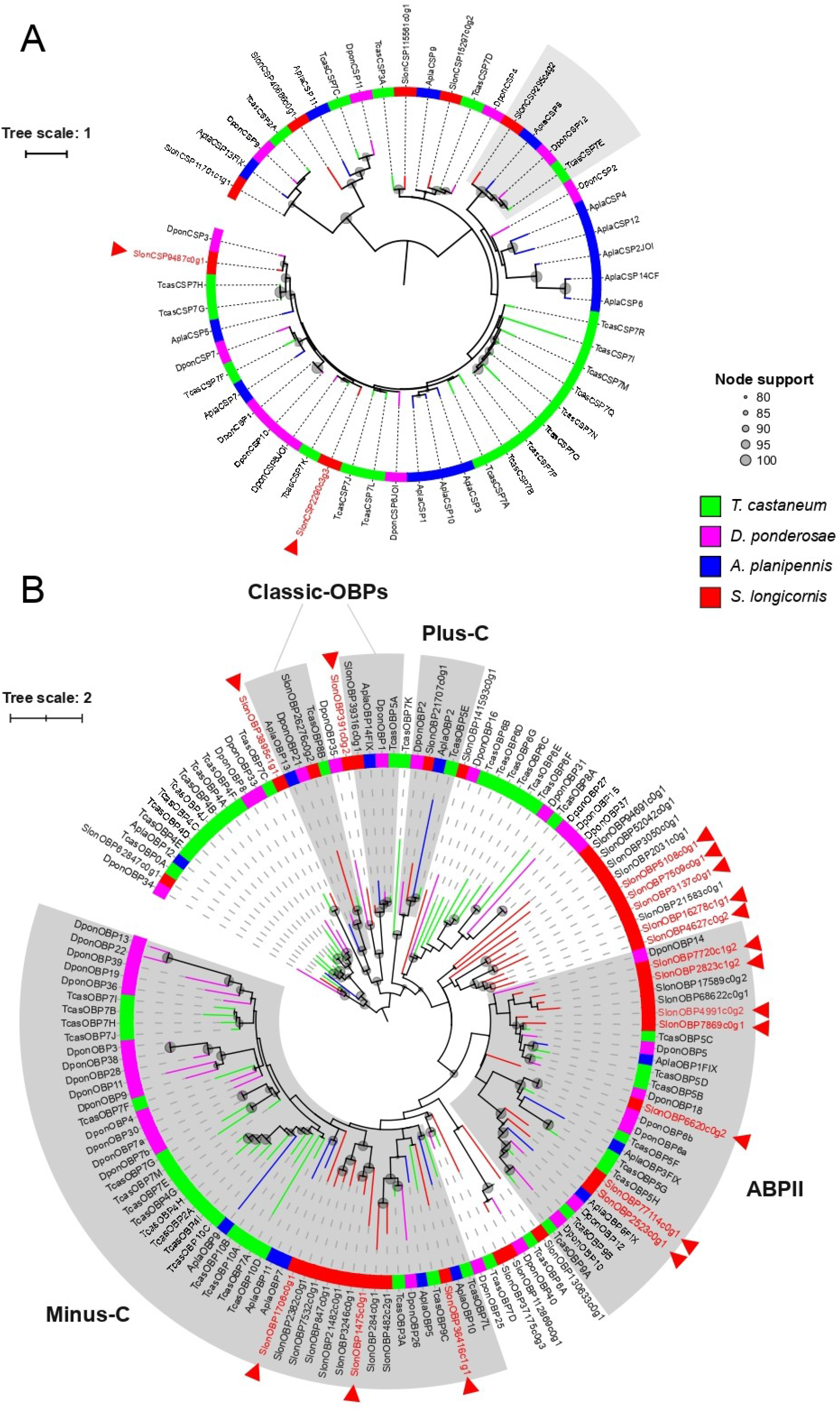
Maximum likelihood phylogenetic trees of (A) chemosensory proteins (CSPs) and (B) odorant-binding proteins (OBPs) including sequences of *S. longicornis* and other coleopterans from Andersson et al. (2019). Grey ranges represent the main OBP clades that were described in previous studies. Red triangles represent upregulated genes in the antennae of *S. longicornis*.

A total of 137 OBP sequences were included to explore the OBPs diversity of *S. longicornis*, resulting in a multiple sequence alignment of 110 amino acid positions after trimming. Our results suggest that OBPs in *S. longicornis* are relatively abundant compared to the other species, being the most diverse repertoire of this comparison after *T. castaneum* (fig. 7B). Several OBP candidates of *S. longicornis* clustered together with the OBPs subgroups described for the other species in Andersson et al. (2019) (i.e., classic-OBPs, minus-C, plus-C and antennal binding proteins II (ABPII)). Only 7 out of the 17 OBPs upregulated in antennae correspond to the ABPII clade. Furthermore, two relatively large OBP expansions include the majority of the upregulated OBPs, one in the minus-C clade (with three upregulated in antennae), and the second forming a specific *S. longicornis* OBP lineage of ten genes (with five upregulated in the antennae).

## DISCUSSION

A highly complete transcriptome for the cave-dwelling beetle *Speonomus longicornis* was generated in the present study (supplementary figure S1). Combining the differential gene expression and GO enrichment analyses with a curated annotation pipeline for the chemosensory related genes, we were able to explore in detail the chemosensory gene repertoire of *S. longicornis*. Furthermore, the phylogenetic inferences for each of the chemosensory gene families offered the opportunity to compare the repertoire of genes involved in chemosensation in *S. longicornis* to other beetle species that occupy a wide variety of ecological niches, all of them in epigean habitats.

The differential gene expression (fig. 2A) and GO enrichment analysis (fig. 2B) allowed us to identify upregulated genes in antennae and compare the overall enriched functions in the antennae versus the rest of the body. Therefore this approach allowed us to pinpoint the genes orchestrating chemosensation in *S. longicornis* particularly in antennae, where chemosensory structures - sensilla - are highly concentrated. As expected, olfaction was recovered as the most prominent function in antennae, representing more than half of the enriched terms in the molecular function category (odorant-binding and olfactory reception activities), indicating that ORs and OBPs are playing a major role in how cave beetles receive and process airborne cues. The differential expression analyses detected a large number of upregulated chemosensory genes in the antennae compared to the rest of the body. ORCO is upregulated in antennae, as expected since it is an essential component of the functional heterodimers that facilitate odorant reception combined with other ORs (Stengl and Funk 2013). In addition, only 7 ORs were observed as exclusively expressed in antennae (supplementary table S4), suggesting that this gene family may include genes with high specificity in these appendages. All in all, these results highlight the importance of antennae in odorant perception in this cave beetle; further gene expression studies including additional structures such as mouth appendages would give more detailed insights for the rest of the identified ORs.

Concerning the ORs phylogeny (fig. 3), our results are mostly consistent with what was found in Mitchell et al. (2019) and most importantly, these data allowed us to characterize the distribution and diversity of the identified ORs for *S. longicornis* compared to those reported in the genomes of other species with different ecological strategies. As reported by Mitchell et al. (2019), large gene expansions in several OR groups are highlighted in the polyphagous/generalist species *T. castaneum, O. taurus, A. glabripennis* and *P. serrata,* whereas the oligophagous/specialist *D. ponderosae, L. decemlineata, A. planipennis* and *H. redfordi* exhibit relatively reduced OR repertoires. Therefore, an apparent correlation between the host breadth and the ORs diversity of herbivore Coleoptera is observed, clearly exemplified by the extent and distribution of OR diversity in the wood boring species (i.e., *A. glabripennis*, *D. ponderosae*, *A. planipennis*; Andersson et al. 2019). The insectivorous *C. scrutator* and the scavengers *N. vespilloides* and *S. longicornis* are conceived as polyphagous/generalists but they present a low number of ORs and relatively smaller gene expansions compared to the rest of polyphagous species. Their OR repertoires present a similar distribution across the Coleoptera OR groups, with moderate expansions in groups 1 and 3 and lacking representatives in group 5A. Although we have only studied the expressed ORs of *S. longicornis*, our results suggest a relatively reduced OR repertoire of *S. longicornis* compared to the other species. Its OR repertoire may result from adaptation to the highly specific ecological niche it occupies. Into the deep subterranean, odorant compounds are more homogeneous and less diverse than in surface habitats, basically due to the absence of light and primary production and therefore probably less complex in chemical information. The OR diversity in this cave-dwelling beetle species may be associated with this scenario of higher simplicity in terms of chemical airborne cues.

Regarding gustatory perception (fig. 4), five GRs were significantly enriched in the antennae in *S. longicornis*, indicating a substantial gustatory role in these appendages. This result is consistent with what was found in the GR expression levels of *T. castaneum*, where they also report similar values in the maxillary palps and the antennae (Dippel et al. 2016). Remarkably, no GRs associated with the perception of fructose and other sugars were detected in *S. longicornis* (at least clustering together with functionally annotated genes in *D. melanogaster*) indicating that either *S. longicornis* does not have receptors for these types of carbohydrates, or their evolutionary origin is different from that in other beetles. This could also be the consequence of reduced proportion of carbohydrates available underground and with the low sugar feeding habits of *S. longicornis*, which basically consist of fungi, biofilms and carrion (Delay 1978; Dorigo et al. 2017). Further comparative studies including non-phytophagous epigean beetles would help to test the hypothesis that a lack of sugar receptors is directly associated with a strict subterranean lifestyle.

CO_2_ perception may be crucial for *S. longicornis* to orientate within its habitat and to detect decomposing organic matter in the darkness, the main food source for this species. The GR-CO_2_ sensing complex was characterized in *S. longicornis*. We detected three candidate GRs clustering together with highly conserved CO_2_ receptors of insects (Robertson and Kent 2009), among which only one candidate was significantly expressed in antennae (fig. 4). Our results suggest that CO_2_ perception may not be restricted to a single chemosensory structure, congruent with what was found in *T. castaneum* after comparing different body structures (Dippel et al. 2016). These results in beetles are in contrast to what was found in well studied dipterans. For instance, *D. melanogaster* has only two CO_2_ receptors that form functional heteromers that are significantly enriched in antennae (i.e., DmelGR21a and DmelGR63a; Jones et al. 2007; Kwon et al. 2007), whereas *A. gambiae* has three CO_2_ receptors that are upregulated in the mouthparts (AgamGR22-24; Pitts et al. 2011). Further studies exploring differential gene expression in different body parts will be needed to deepen our understanding on CO_2_ perception in *S. longicornis*.

The GR repertoire of *S. longicornis* was small and showed similar to that observed in the oligophagous *A. planipennis* (ash tree specialist). Our findings on the gustatory perception of this cavernicolous beetle may reflect the poor diversity of gustatory substances in the hypogean habitat compared to surface environments. Further investigation with genomic data will confirm the lack of sugar receptors in this subterranean species and will help to assess the extent of its gustatory repertoire.

The IRs/IGluRs gene family can be classified in two subfamilies: the ancestral ionotropic glutamate receptors (iGluRs) and the recently described subfamily of ionotropic receptors (IRs), which include divergent ligand-binding domains that lack their characteristic glutamate-interacting residues (Benton et al. 2009; Croset et al. 2010). Contrary to the specific role of iGluRs in synaptic communication, IRs have more diverse roles which in insects are often related to chemoreception (Rytz et al. 2013; Koh et al. 2014). While the most conserved IRs (e.g., IR8a and IR25a) act as co-receptors conferring multiple odor-evoked electrophysiological responses, more recently some insect IRs have been found to mediate specific stimulus forming heterodimers with more selectively expressed IR subunits (Abuin et al. 2011; Abuin et al. 2019). For instance, in *D. melanogaster*, the highly conserved coreceptors IR93a and IR25a are coexpressed with IR21a, mediating physiological and behavioral responses to low temperatures (Ni et al. 2016; Knecht et al. 2017). In *S. longicornis* we found overexpression in the antennae of the candidate genes clustering together to *D. melanogaster* IR93a and IR25a, while the candidate IR21a (SlonIR2299c0g) was significantly underexpressed in these appendages (fig. 5C). By contrast, higher expression levels for IR21a of *D. melanogaster* were found in the antennae (Sánchez-Alcañiz et al. 2018) and the same was found for other Coleoptera (Dippel et al. 2016; Bin et al. 2017). However, we did not explore differential gene expression in different structures from the body, which could explain the differences in the observed results.

Remarkably, our results suggest a putative duplication in the two genes annotated as IR25a, despite being a highly conserved gene with a single copy in virtually all protostomes (fig. 5B). This duplication has been only reported for the limpet *L. gigantea* and for two parasitoid wasp species: *N. vitripennis and M. mediator* (Croset et al. 2010; Wang et al. 2015). The inferred phylogeny suggested different origins for the observed duplication in the IR25a candidates: while a recent and lineage-specific duplication has been observed in *N. vitripennis* and *M. mediator,* where both copies of IR25a were retrieved as sister to each other, the candidate duplication in *S. longicornis* seems to be older and may represent an ancestral duplication in Coleoptera that was retained in this cave beetle. Our results represent the first report of a gene duplication observed in this highly conserved gene in Coleoptera, which may indicate that the evolutionary history of IR25a and its role in chemoreception may be more complex than originally considered across arthropods.

Cave beetles inhabit a medium where air tends to be still and the ambient temperature and humidity fluctuate only by tiny amounts over long periods, and therefore a good thermal detection may have a selective advantage. Physiological experiments on a closer relative species (i.e., the cave-dwelling *Speophyes lucidulus,* Leoididae, Cholevinae) revealed an extreme sensitivity to small changes in temperature incurred by antennal receptors (Corbière-Tichané and Loftus 1983) that may be mediated by some of the inferred candidate IRs. Consequently, other relevant IRs for *S. longicornis* may be those potentially related to humidity sensing. The functionally characterized IR40a and IR68a in *D. melanogaster* have been seen to be coexpressed with IR93a and IR25a in specialized sensory neurons of the antennae performing hygrosensory responses (Enjin 2017; Knecht et al. 2017). Through the phylogenetic analysis we identified the hygroreceptor candidates (IR40a and IR68a) for *S. longicornis* (fig. 5C), although we did not find significant differences in the expression values between antennae and the rest of the body. The rest of candidate IRs annotated in *S. longicornis* (i.e., IR8a, IR76b, IR75a, IR64a, 100a and IR60a; see fig. 5C and supplementary table S2) have been shown to be potentially involved in taste and odor transduction *in D. melanogaster*, suggesting candidate odorant and gustatory roles in *S. longicornis*.

Regarding the genes encoding sensory neuron membrane proteins (SNMPs/CD36s gene family), their apparent conservation between phylogenetically distant species may suggest conserved protein functions. Previous studies on insects classified these genes fall into three major groups (Nichols and Vogt 2008; Vogt et al. 2009). Little is known about the highly conserved group 1 formed by the epithelial membrane proteins (emp), while group 2 includes the previously characterized *D. melanogaster* genes croquemort (Crq), ninaD, santa maria (SM), and peste (Pes). These proteins have a notably similar function to the one described for the CD36 family and its vertebrate relatives (e.g., small ligand transport or cytoadhesion). The group 3 proteins, termed as SNMPs, appear to be quite distinct from the others in their apparent association with chemosensory organs, and yet perhaps similar to the suggested CD36 role in vertebrate taste transduction (SNMP1 and SNMP2). Moreover, SNMP3 was more recently described exclusively for the divergent SNMPs found in Coleoptera (Dippel et al. 2016). The upregulated genes in antennae correspond to groups 2 and 3, indicating that not only SNMPs with putative chemosensory roles are enriched in antennae (fig. 6). Furthermore, SNMP1 has been suggested to play a key role in pheromone perception in *D. melanogaster* and moths (Benton et al. 2007; Pregitzer et al. 2014; Gomez-Diaz et al. 2016). As in *A. planipennis,* a single gene clustering within SNMP1 was found in *S. longicornis,* which was upregulated in antennae, suggesting that pheromone detection is mediated by SNMPs (among other potential genes) in antennae.

Our results suggested that *S. longicornis* does not exhibit species-specific CSPs expansions (fig. 7A). Four of the seven CSPs identified in *S. longicornis* were not enriched in antennae or in the body, similar to what was found in *T. castaneum,* where the majority of CSPs were not found to be significantly enriched at any of the tested structures (Dippel et al. 2014). Further information about structure-specific expression in other Coleoptera would be needed to test the apparently versatile roles of CSPs in chemosensory transduction.

Regarding OBPs, the GO enrichment analysis highlighted odorant-binding functions vastly enriched in antennae (fig. 2B). In addition, the differential gene expression analyses identified a high number of upregulated genes (17 in antennae; fig. 2A). The large number of OBPs annotated in *S. longicornis* suggests a relatively diverse repertoire with species-specific gene duplications and expansions; this may indicate an important role of these proteins in odorant perception in this subterranean beetle. Notably, less than half of the upregulated OBPs in antennae clustered together with the previously described “antennal binding proteins II” (ABPII) in Vieira and Rozas (2011) (fig. 7B). These results indicate that although ABPII were described as OBPs typically enriched in antennae, some genes of this clade may also be differentially expressed in some other body structures, as also found for *T. castaneum* (Dippel et al. 2014). The rest of the upregulated OBPs in both conditions clustered together with the different OBP groups described in previous studies (Vieira and Rozas 2011; Dippel et al. 2014; Andersson et al. 2019). Further research including closer relatives to this cave species with different ecological preferences will allow us to test the hypothesis that subterranean specialization has directed the olfactory and other chemosensory capabilities in Coleoptera.

## CONCLUSION

In this piece of work, we characterized for the first time the chemosensory gene repertoire of an obligate subterranean species, the cave-dwelling coleopteran *S. longicornis*. We found relatively diminished odorant and gustatory repertoires compared to epigean polyphagous coleopterans and more similar to that of those considered specialists based on their feeding habits. Considering the selective pressures of the niche that *S. longicornis* occupies (i.e., limited resources, poor diversity and heterogeneous distribution of food, or low complexity of airborne cues, among others), an optimized chemosensory repertoire in terms of diversity may result from its adaptation to the deep subterranean, reducing the associated energetic costs of maintaining a highly diverse gene repertoire. In this obligate cave-dwelling beetle, we identified some putative gene losses (e.g., sugar gustatory receptors and IR41a) and a relatively reduced diversity of its gustatory and odorant gene repertoires compared to epigean coleopterans. Furthermore, several gene duplications and expansions were observed, epitomized by a duplication of the gene IR25a, which might potentially have facilitated adaptation to subterranean conditions in this cave beetle. Our study thus paves the road towards a better understanding of how subterranean animals perceive their particular environment.

## DATA AVAILABILITY

Raw reads have been deposited in the National Center for Biotechnology Information (NCBI) (BioProject accession number PRJNA667243, BioSample accession number SAMN16362632). Assembled sequences, alignments and custom scripts have been deposited in github (https://github.com/MetazoaPhylogenomicsLab).

## ACKNOWLEDGEMENTS

This work was supported by the Ministerio de Economía y Competitividad and the Ministerio de Ciencia of Spain (CGL2016-76705-P to IR, PID2019-108824GA-I00 to RF, and CGL2016-75255 and PID2019-103947GB to JR). PB and PE were supported by an FPI grant (Ministerio de Economía y Competitividad BES-2017-081050 and BES-2017-081740, respectively). RF was supported by a Marie Sklodowska-Curie grant (747607) and a Ramón y Cajal fellowship (Ministerio de Economía y Competitividad, RyC2017-22492). C. Vanderbergh and E. Ruzzier kindly provided the images for figures 1A and 1B respectively; the latter was obtained at the Electron Microscope Service of the Institute of Marine Sciences (ICM) of Barcelona, Spain. We are grateful for the support received by the CNAG Sequencing facility, which performed the experiments related to the sequencing with great care. This paper is dedicated to the memory of our wonderful colleague, Ignacio Ribera, who worked intensively on this project until his very last days and sadly passed away recently.

## REFERENCES

Abuin L, Bargeton B, Ulbrich MH, Isacoff EY, Kellenberger S, Benton R. 2011. Functional architecture of olfactory ionotropic glutamate receptors. Neuron 69:44–60.

Abuin L, Prieto-Godino LL, Pan H, Gutierrez C, Huang L, Jin R, Benton R. 2019. In vivo assembly and trafficking of olfactory Ionotropic Receptors. BMC Biology 17:34.

Almudi I, Vizueta J, Wyatt CDR, de Mendoza A, Marlétaz F, Firbas PN, Feuda R, Masiero G, Medina P, Alcaina-Caro A, et al. 2020. Genomic adaptations to aquatic and aerial life in mayflies and the origin of insect wings. Nature Communications 11:2631.

Al-Shahrour F, Minguez P, Tárraga J, Medina I, Alloza E, Montaner D, Dopazo J. 2007. FatiGO +: a functional profiling tool for genomic data. Integration of functional annotation, regulatory motifs and interaction data with microarray experiments. Nucleic Acids Research 35:W91–W96.

Andersson MN, Keeling CI, Mitchell RF. 2019. Genomic content of chemosensory genes correlates with host range in wood-boring beetles (Dendroctonus ponderosae, Agrilus planipennis, and Anoplophora glabripennis). BMC Genomics 20:690.

Anholt RRH. 2020. Chemosensation and Evolution of Drosophila Host Plant Selection. iScience 23:100799.

Benjamini Y, Hochberg Y. 1995. Controlling the False Discovery Rate: A Practical and Powerful Approach to Multiple Testing. Journal of the Royal Statistical Society: Series B (Methodological) 57:289–300.

Benton R. 2015. Multigene Family Evolution: Perspectives from Insect Chemoreceptors. Trends in Ecology & Evolution 30:590–600.

Benton R, Vannice KS, Gomez-Diaz C, Vosshall LB. 2009. Variant Ionotropic Glutamate Receptors as Chemosensory Receptors in Drosophila. Cell 136:149–162.

Benton R, Vannice KS, Vosshall LB. 2007. An essential role for a CD36-related receptor in pheromone detection in Drosophila. Nature 450:289–293.

Bin S-Y, Qu M-Q, Li K-M, Peng Z-Q, Wu Z-Z, Lin J-T. 2017. Antennal and abdominal transcriptomes reveal chemosensory gene families in the coconut hispine beetle, Brontispa longissima. Scientific Reports 7:2809.

Bolger AM, Lohse M, Usadel B. 2014. Trimmomatic: a flexible trimmer for Illumina sequence data. Bioinformatics 30:2114–2120.

Capella-Gutiérrez S, Silla-Martínez JM, Gabaldón T. 2009. trimAl: a tool for automated alignment trimming in large-scale phylogenetic analyses. Bioinformatics 25:1972–1973.

Cartwright RA, Schwartz RS, Merry AL, Howell MM. 2017. The importance of selection in the evolution of blindness in cavefish. BMC Evolutionary Biology 17:45.

Cheon S, Zhang J, Park C. 2020. Is Phylotranscriptomics as Reliable as Phylogenomics? Molecular Biology and Evolution.

Cieslak A, Fresneda J, Ribera I. 2014. Life-history specialization was not an evolutionary dead-end in Pyrenean cave beetles. Proceedings of the Royal Society B: Biological Sciences 281:20132978.

Clyne PJ, Warr CG, Freeman MR, Lessing D, Kim J, Carlson JR. 1999. A novel family of divergent seven-transmembrane proteins: candidate odorant receptors in Drosophila. Neuron 22:327–338.

Cock PJA, Antao T, Chang JT, Chapman BA, Cox CJ, Dalke A, Friedberg I, Hamelryck T, Kauff F, Wilczynski B, et al. 2009. Biopython: freely available Python tools for computational molecular biology and bioinformatics. Bioinformatics 25:1422–1423.

Corbière-Tichané G, Loftus R. 1983. Antennal thermal receptors of the cave beetle,Speophyes lucidulus Delar. Journal of Comparative Physiology 153:343–351.

Croset V, Rytz R, Cummins SF, Budd A, Brawand D, Kaessmann H, Gibson TJ, Benton R. 2010. Ancient protostome origin of chemosensory ionotropic glutamate receptors and the evolution of insect taste and olfaction. PLOS Genetics 6:e1001064.

Croset V, Schleyer M, Arguello JR, Gerber B, Benton R. 2016. A molecular and neuronal basis for amino acid sensing in the Drosophila larva. Scientific Reports 6:34871.

Delay B. 1978. Milieu souterrain et ecophysiologie de la reproduction et du développement des coléoptères Bathysciinae hypoges. Mémoires de Biospéologie 5:1–349.

Deleurance S. 1963. Recherches sur les coléoptères troglobies de la sous-famille des Bathysciinae. Annales des Sciences Naturelles (Paris) Zoologie 1:1–172.

Dippel S, Kollmann M, Oberhofer G, Montino A, Knoll C, Krala M, Rexer K-H, Frank S, Kumpf R, Schachtner J, et al. 2016. Morphological and Transcriptomic Analysis of a Beetle Chemosensory System Reveals a Gnathal Olfactory Center. BMC Biology 14:90.

Dippel S, Oberhofer G, Kahnt J, Gerischer L, Opitz L, Schachtner J, Stanke M, Schütz S, Wimmer EA, Angeli S. 2014. Tissue-specific transcriptomics, chromosomal localization, and phylogeny of chemosensory and odorant binding proteins from the red flour beetle Tribolium castaneum reveal subgroup specificities for olfaction or more general functions. BMC Genomics 15:1141.

Dorigo L, Squartini A, Toniello V, Dreon AL, Pamio A, Concina G, Simonutti V, Ruzzier E, Perreau M, Engel AS, et al. 2017. Cave hygropetric beetles and their feeding behaviour, a comparative study of Cansiliella servadeii AND Hadesia asamo (Coleoptera, Leiodidae, Cholevinae, Leptodirini). Acta Carsologica 46.

Enjin A. 2017. Humidity sensing in insects-from ecology to neural processing. Current Opinion in Insect Science 24:1–6.

Eyun S-I, Soh HY, Posavi M, Munro JB, Hughes DST, Murali SC, Qu J, Dugan S, Lee SL, Chao H, et al. 2017. Evolutionary History of Chemosensory-Related Gene Families across the Arthropoda. Molecular Biology and Evolution 34:1838–1862.

Fernández R, Gabaldón T. 2020. Gene gain and loss across the metazoan tree of life. Nature Ecology and Evolution 4:524–533.

Gao Q, Chess A. 1999. Identification of candidate Drosophila olfactory receptors from genomic DNA sequence. Genomics 60:31–39.

Glaçon S. 1953. The evolutive cycle of a cave coleopter Speonomus longicornis Saulcy. Comptes Rendus Hebdomadaires des Séances de l’Académie des Sciences. 236:2443–2445.

Gomez-Diaz C, Bargeton B, Abuin L, Bukar N, Reina JH, Bartoi T, Graf M, Ong H, Ulbrich MH, Masson J-F, et al. 2016. A CD36 ectodomain mediates insect pheromone detection via a putative tunnelling mechanism. Nature Communications 7.

Grabherr MG, Haas BJ, Yassour M, Levin JZ, Thompson DA, Amit I, Adiconis X, Fan L, Raychowdhury R, Zeng Q, et al. 2011. Full-length transcriptome assembly from RNA-Seq data without a reference genome. Nature Biotechnology 29:644–652.

Grimaldi D, Engel MS. 2005. Evolution of the Insects. Cambridge University Press

Haas BJ, Papanicolaou A, Yassour M, Grabherr M, Blood PD, Bowden J, Couger MB, Eccles D, Li B, Lieber M, et al. 2013. De novo transcript sequence reconstruction from RNA-seq using the Trinity platform for reference generation and analysis. Nature Protocols 8:1494–1512.

Hoang DT, Chernomor O, von Haeseler A, Minh BQ, Vinh LS. 2018. UFBoot2: Improving the Ultrafast Bootstrap Approximation. Molecular Biology and Evolution 35:518–522.

Hörnschemeyer T, Bond J, Young PG. 2013. Analysis of the functional morphology of mouthparts of the beetle Priacma serrata, and a discussion of possible food sources. Journal of Insect Science 13:126.

Huerta-Cepas J, Forslund K, Coelho LP, Szklarczyk D, Jensen LJ, von Mering C, Bork P. 2017. Fast Genome-Wide Functional Annotation through Orthology Assignment by eggNOG-Mapper. Molecular Biology and Evolution 34:2115–2122.

Jeannel R. 1924. Monographie des Bathysciinae. Archives de Zoologie Expérimentale et Générale (Paris) 63:1–436.

Jones WD, Cayirlioglu P, Kadow IG, Vosshall LB. 2007. Two chemosensory receptors together mediate carbon dioxide detection in Drosophila. Nature 445:86–90.

Joseph RM, Carlson JR. 2015. Drosophila Chemoreceptors: A Molecular Interface Between the Chemical World and the Brain. Trends in Genetics 31:683–695.

Knecht ZA, Silbering AF, Cruz J, Yang L, Croset V, Benton R, Garrity PA. 2017. Ionotropic Receptor-dependent moist and dry cells control hygrosensation in Drosophila. eLife 6. Available from: http://dx.doi.org/10.7554/elife.26654

Koh T-W, He Z, Gorur-Shandilya S, Menuz K, Larter NK, Stewart S, Carlson JR. 2014. The Drosophila IR20a clade of ionotropic receptors are candidate taste and pheromone receptors. Neuron 83:850–865.

Kwon JY, Dahanukar A, Weiss LA, Carlson JR. 2007. The molecular basis of CO2 reception in Drosophila. Proceedings of the National Academy of Sciences of the United States of America 104:3574–3578.

Laetsch DR, Blaxter ML. 2017. BlobTools: Interrogation of genome assemblies. F1000Research 6:1287.

Langmead B, Salzberg SL. 2012. Fast gapped-read alignment with Bowtie 2. Nature Methods 9:357–359.

Letunic I, Bork P. 2019. Interactive Tree Of Life (iTOL) v4: recent updates and new developments. Nucleic Acids Research 47:W256–W259.

Luo X-Z, Antunes-Carvalho C, Wipfler B, Ribera I, Beutel RG. 2019. The cephalic morphology of the troglobiontic cholevine species Troglocharinus ferreri (Coleoptera, Leiodidae). Journal of Morphology 280:1207–1221.

McKenna DD, Shin S, Ahrens D, Balke M, Beza-Beza C, Clarke DJ, Donath A, Escalona HE, Friedrich F, Letsch H, et al. 2019. The evolution and genomic basis of beetle diversity. Proceedings of the National Academy of Sciences of the United States of America 116:24729–24737.

Mirarab S, Nguyen N, Guo S, Wang L-S, Kim J, Warnow T. 2015. PASTA: Ultra-Large Multiple Sequence Alignment for Nucleotide and Amino-Acid Sequences. Journal of Computational Biology 22:377–386.

Missbach C, Dweck HKM, Vogel H, Vilcinskas A, Stensmyr MC, Hansson BS, Grosse-Wilde E. 2014. Evolution of insect olfactory receptors. eLife 3.

Missbach C, Vogel H, Hansson BS, Groβe-Wilde E. 2015. Identification of Odorant Binding Proteins and Chemosensory Proteins in Antennal Transcriptomes of the Jumping Bristletail Lepismachilis y-signata and the Firebrat Thermobia domestica: Evidence for an Independent OBP-OR Origin. Chemical Senses 40:615–626.

Mitchell RF, Schneider TM, Schwartz AM, Andersson MN, McKenna DD. 2019. The diversity and evolution of odorant receptors in beetles (Coleoptera). Insect Molecular Biology 29:77–91.

Nei M, Niimura Y, Nozawa M. 2008. The evolution of animal chemosensory receptor gene repertoires: roles of chance and necessity. Nature Reviews Genetics 9:951–963.

Nei M, Rooney AP. 2005. Concerted and Birth-and-Death Evolution of Multigene Families. Annual Review of Genetics 39:121–152.

Nguyen L-T, Schmidt HA, von Haeseler A, Minh BQ. 2015. IQ-TREE: a fast and effective stochastic algorithm for estimating maximum-likelihood phylogenies. Molecular Biology and Evolution 32:268–274.

Nichols Z, Vogt RG. 2008. The SNMP/CD36 gene family in Diptera, Hymenoptera and Coleoptera: Drosophila melanogaster, D. pseudoobscura, Anopheles gambiae, Aedes aegypti, Apis mellifera, and Tribolium castaneum. Insect Biochemistry and Molecular Biology 38:398–415.

Ni L, Klein M, Svec KV, Budelli G, Chang EC, Ferrer AJ, Benton R, Samuel AD, Garrity PA. 2016. The Ionotropic Receptors IR21a and IR25a mediate cool sensing in Drosophila. eLife 5.

Nishimura O, Hara Y, Kuraku S. 2017. gVolante for standardizing completeness assessment of genome and transcriptome assemblies. Bioinformatics 33:3635–3637.

Pallarés S, Ribera I, Montes A, Millán A, Rizzo V, Comas J, Sánchez-Fernández D. 2018. Limited thermal acclimation capacity in cave beetles. ARPHA Conference Abstracts 1.

Parzefall J. 2001. A review of morphological and behavioural changes in the cave molly, Poecilia mexicana, from Tabasco, Mexico. The biology of hypogean fishes:263–275.

Patro R, Duggal G, Love MI, Irizarry RA, Kingsford C. 2017. Salmon provides fast and bias-aware quantification of transcript expression. Nature Methods 14:417–419.

Pelosi P, Iovinella I, Felicioli A, Dani FR. 2014. Soluble proteins of chemical communication: an overview across arthropods. Frontiers in Physiology 5:320.

Pelosi P, Iovinella I, Zhu J, Wang G, Dani FR. 2018. Beyond chemoreception: diverse tasks of soluble olfactory proteins in insects. Biological reviews of the Cambridge Philosophical Society 93:184–200.

Picelli S, Björklund ÅK, Faridani OR, Sagasser S, Winberg G, Sandberg R. 2013. Smart-seq2 for sensitive full-length transcriptome profiling in single cells. Nature Methods 10:1096–1098.

Pipan T, Culver DC. 2012. Convergence and divergence in the subterranean realm: a reassessment. Biological Journal of the Linnean Society 107:1–14.

Pitts RJ, Rinker DC, Jones PL, Rokas A, Zwiebel LJ. 2011. Transcriptome profiling of chemosensory appendages in the malaria vector Anopheles gambiae reveals tissue- and sex-specific signatures of odor coding. BMC Genomics 12:271.

Pregitzer P, Greschista M, Breer H, Krieger J. 2014. The sensory neurone membrane protein SNMP1 contributes to the sensitivity of a pheromone detection system. Insect Molecular Biology 23:733–742.

Price MN, Dehal PS, Arkin AP. 2010. FastTree 2--approximately maximum-likelihood trees for large alignments. PLoS One 5:e9490.

Ribera I, Fresneda J, Bucur R, Izquierdo A, Vogler AP, Salgado JM, Cieslak A. 2010. Ancient origin of a Western Mediterranean radiation of subterranean beetles. BMC Evolutionary Biology 10:29.

Rizzo V, Sánchez-Fernández D, Fresneda J, Cieslak A, Ribera I. 2015. Lack of evolutionary adjustment to ambient temperature in highly specialized cave beetles. BMC Evolutionary Biology 15:10.

Robertson HM. 2019. Molecular Evolution of the Major Arthropod Chemoreceptor Gene Families. Annual Review of Entomology 64:227–242.

Robertson HM, Kent LB. 2009. Evolution of the gene lineage encoding the carbon dioxide receptor in insects. Journal of Insect Science 9:19.

Robertson HM, Warr CG, Carlson JR. 2003. Molecular evolution of the insect chemoreceptor gene superfamily in Drosophila melanogaster. Proceedings of the National Academy of Sciences of the United States of America 100 Suppl 2:14537–14542.

Robinson MD, McCarthy DJ, Smyth GK. 2010. edgeR: a Bioconductor package for differential expression analysis of digital gene expression data. Bioinformatics 26:139–140.

Robinson MD, Oshlack A. 2010. A scaling normalization method for differential expression analysis of RNA-seq data. Genome Biology 11:R25.

Roys C. 1954. Olfactory nerve potentials a direct measure of chemoreception in insects. Annals of the New York Academy of Sciences 58:250–255.

Rytz R, Croset V, Benton R. 2013. Ionotropic receptors (IRs): chemosensory ionotropic glutamate receptors in Drosophila and beyond. Insect Biochemistry and Molecular Biology 43:888–897.

Sánchez-Alcañiz JA, Silbering AF, Croset V, Zappia G, Sivasubramaniam AK, Abuin L, Sahai SY, Münch D, Steck K, Auer TO, et al. 2018. An expression atlas of variant ionotropic glutamate receptors identifies a molecular basis of carbonation sensing. Nature Communications 9:4252.

Sánchez-Gracia A, Vieira FG, Rozas J. 2009. Molecular evolution of the major chemosensory gene families in insects. Heredity 103:208–216.

Simão FA, Waterhouse RM, Ioannidis P, Kriventseva EV, Zdobnov EM. 2015. BUSCO: assessing genome assembly and annotation completeness with single-copy orthologs. Bioinformatics 31:3210–3212.

Stengl M, Funk NW. 2013. The role of the coreceptor Orco in insect olfactory transduction. Journal of Comparative Physiology A: Neuroethology, Sensory, Neural, and Behavioral Physiology 199:897–909.

Stocker RF. 1994. The organization of the chemosensory system in Drosophila melanogaster: a rewiew. Cell and Tissue Research 275:3–26.

Stürckow B. 1970. Responses of Olfactory and Gustatory Receptor Cells in Insects. Advances in Chemoreception:107–159.

Supek F, Bošnjak M, Škunca N, Šmuc T. 2011. REVIGO summarizes and visualizes long lists of gene ontology terms. PLoS One 6:e21800.

Thoma M, Missbach C, Jordan MD, Grosse-Wilde E, Newcomb RD, Hansson BS. 2019. Transcriptome Surveys in Silverfish Suggest a Multistep Origin of the Insect Odorant Receptor Gene Family. Frontiers in Ecology and Evolution 7:281.

Turk S, Sket B, Sarbu ?erban. 1996. Comparison between some epigean and hypogean populations of Asellus aquaticus (Crustacea: Isopoda: Asellidae). Hydrobiologia 337:161–170.

Vieira FG, Rozas J. 2011. Comparative Genomics of the Odorant-Binding and Chemosensory Protein Gene Families across the Arthropoda: Origin and Evolutionary History of the Chemosensory System. Genome Biology and Evolution 3:476–490.

Vieira FG, Sánchez-Gracia A, Rozas J. 2007. Comparative genomic analysis of the odorant-binding protein family in 12 Drosophila genomes: purifying selection and birth-and-death evolution. Genome Biology 8:R235.

Vizueta J, Frías-López C, Macías-Hernández N, Arnedo MA, Sánchez-Gracia A, Rozas J. 2016. Evolution of chemosensory gene families in arthropods: Insight from the first inclusive comparative transcriptome analysis across spider appendages. Genome Biology and Evolution:evw296.

Vizueta J, Sánchez-Gracia A, Rozas J. 2020. BITACORA: A comprehensive tool for the identification and annotation of gene families in genome assemblies. Molecular Ecology Resources.

Vogt RG, Miller NE, Litvack R, Fandino RA, Sparks J, Staples J, Friedman R, Dickens JC. 2009. The insect SNMP gene family. Insect Biochemistry and Molecular Biology 39:448–456.

Vosshall LB, Amrein H, Morozov PS, Rzhetsky A, Axel R. 1999. A spatial map of olfactory receptor expression in the Drosophila antenna. Cell 96:725–736.

Vosshall LB, Stocker RF. 2007. Molecular architecture of smell and taste in Drosophila. Annual Review of Neuroscience 30:505–533.

Wang D, Pentzold S, Kunert M, Groth M, Brandt W, Pasteels JM, Boland W, Burse A. 2018. A subset of chemosensory genes differs between two populations of a specialized leaf beetle after host plant shift. Ecology and Evolution 8:8055–8075.

Wang H-C, Minh BQ, Susko E, Roger AJ. 2018. Modeling Site Heterogeneity with Posterior Mean Site Frequency Profiles Accelerates Accurate Phylogenomic Estimation. Systematic Biology 67:216–235.

Wang S-N, Peng Y, Lu Z-Y, Dhiloo KH, Gu S-H, Li R-J, Zhou J-J, Zhang Y-J, Guo Y-Y. 2015. Identification and Expression Analysis of Putative Chemosensory Receptor Genes in Microplitis mediator by Antennal Transcriptome Screening. International Journal of Biological Sciences 11:737–751.

Yamamoto Y, Byerly MS, Jackman WR, Jeffery WR. 2009. Pleiotropic functions of embryonic sonic hedgehog expression link jaw and taste bud amplification with eye loss during cavefish evolution. Developmental Biology 330:200–211.

Yang J, Chen X, Bai J, Fang D, Qiu Y, Jiang W, Yuan H, Bian C, Lu J, He S, et al. 2016. The Sinocyclocheilus cavefish genome provides insights into cave adaptation. BMC Biology 14:1.

