## supplementary figure S1 for "Smelling in the dark: phylogenomic insights on the chemosensory system of a subterranean beetle"

**Supplementary figure S1.** Summary of raw data data and read yield per library, basic assembly statistics, assessment of completeness (BUSCO) and assessment of contamination (Blobtools).

### Libraries and RNA-seq information

| Sample code | Condition | Reads | Yield seq Gb | Trimmed Illumina Gb |
|---|---|---|---|---|
| Sl_a_rep1 | antennae | 67.047.081 | 11.308 | 10.188 |
| Sl_a_rep2 | antennae | 55.080.352 | 9.254 | 8.371 |
| Sl_a_rep3 | antennae | 56.297.588 | 9.399 | 8.555 |
| Sl_b_rep1 | body w/o antennae | 74.470.767 | 11.32 | 11.319 |
| Sl_b_rep2 | body w/o antennae | 81.147.359 | 12.334 | 12.331 |
| Sl_b_rep3 | body w/o antennae | 77.169.906 | 13.044 | 11.724 |

### Basic assembly statistics

**Total Trinity 'genes':** 177711
**Total transcripts:** 245131
**Percent GC:** 38.8

Based on all transcripts contigs:

**N50:** 2184
**Median conting length:** 394
**Average contig length:** 954.84
**Total assembled bases:** 235041111

Based on only longuest isoform per 'gene'

**N50:** 957
**Median contig length:** 326
**Average contig length:** 625.35
**Total assembled bases:** 111132230

### ExN50 statistic graph

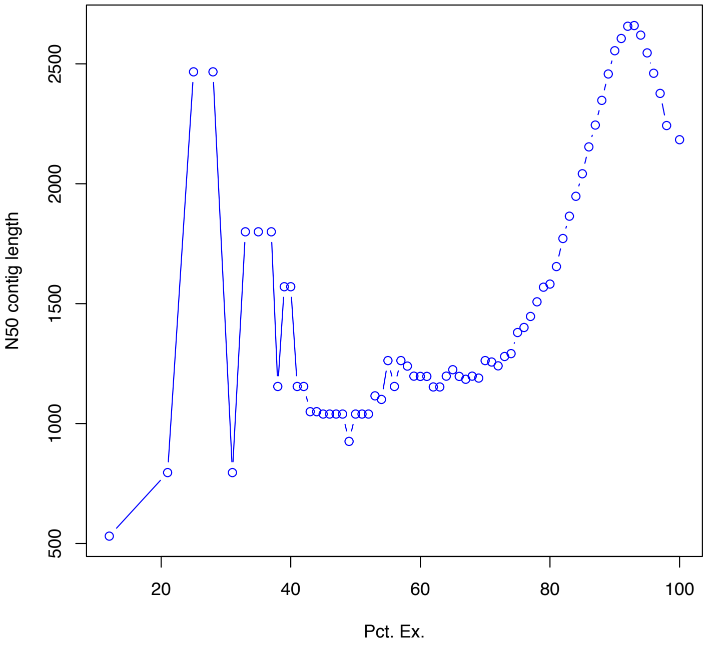


### BUSCO summary:

- Lineage dataset: arthropoda_odb9
- Run mode: transcriptome

- Hits Percentage: C:99.0%[S:20.1%,D:78.9%],F:0.5%,M:0.5%,n:1066
- Hits counts:

| | |
|---|---|
| 1055 | Complete BUSCOs (C) |
| 214 | Complete and single-copy BUSCOs (S) |
| 841 | Complete and duplicated BUSCOs (D) |
| 5 | Fragmented BUSCOs (F) |
| 6 | Missing BUSCOs (M) |
| 1066 | Total BUSCO groups searched |

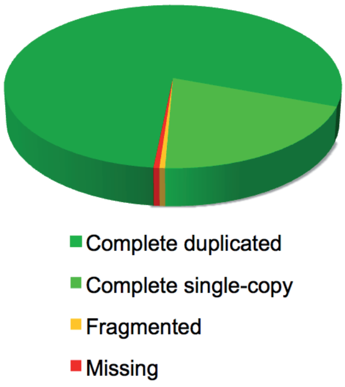


### Blobtools summary:

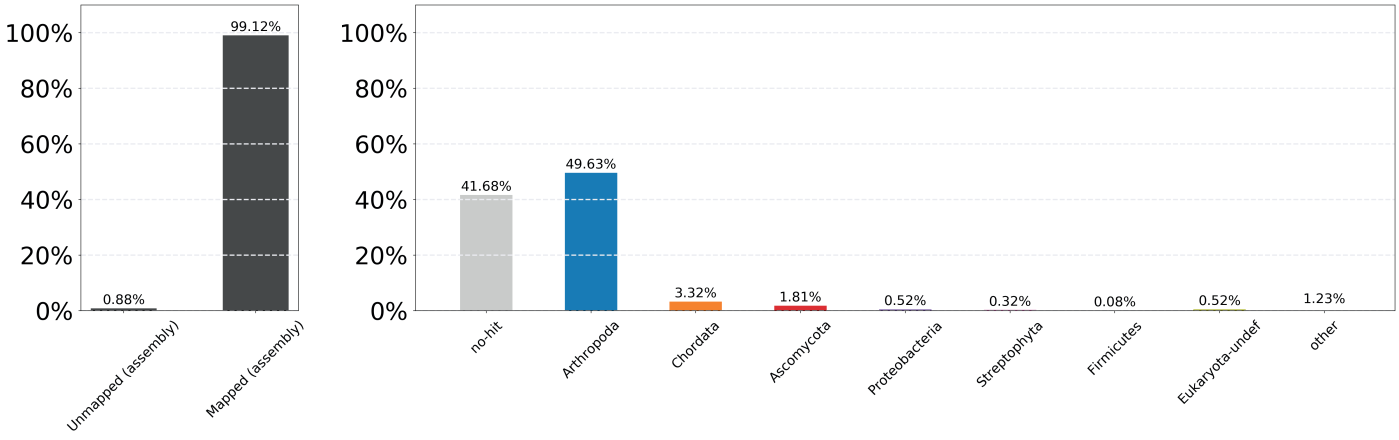
