## supplementary figure S2 for "Smelling in the dark: phylogenomic insights on the chemosensory system of a subterranean beetle"

**Supplementary figure S2.** Replicate correlations and differential gene expression counts between conditions and replicates. Antennae replicates are coded as Sla and the rest of the body replicates as Slb.

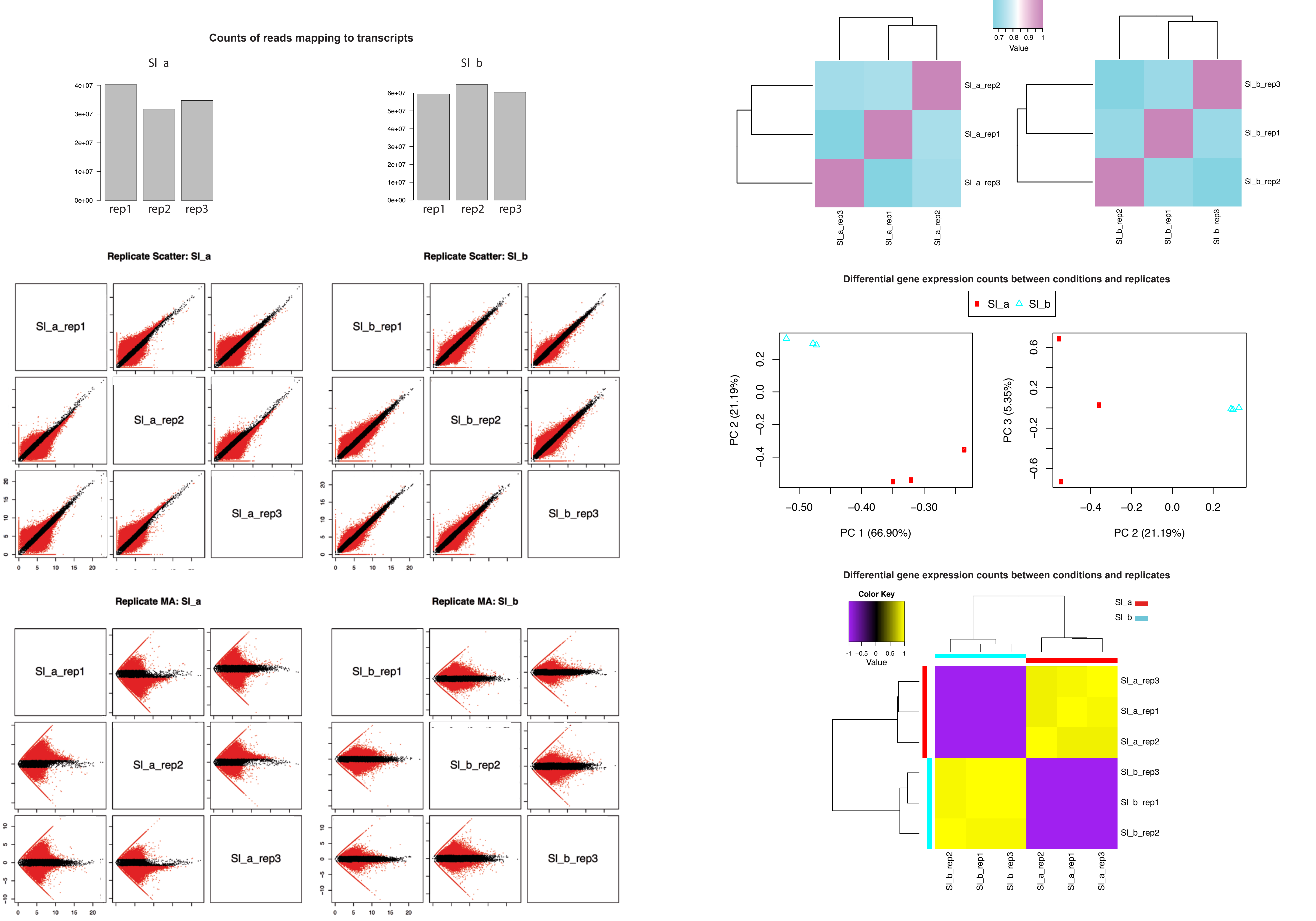
