## supplementary figure S3 for "Smelling in the dark: phylogenomic insights on the chemosensory system of a subterranean beetle"

**Supplementary figure S3.** Heatmap of all differentially expressed genes between antennae (Sla) and the rest of the body (Slb) of *S. longicornis*.

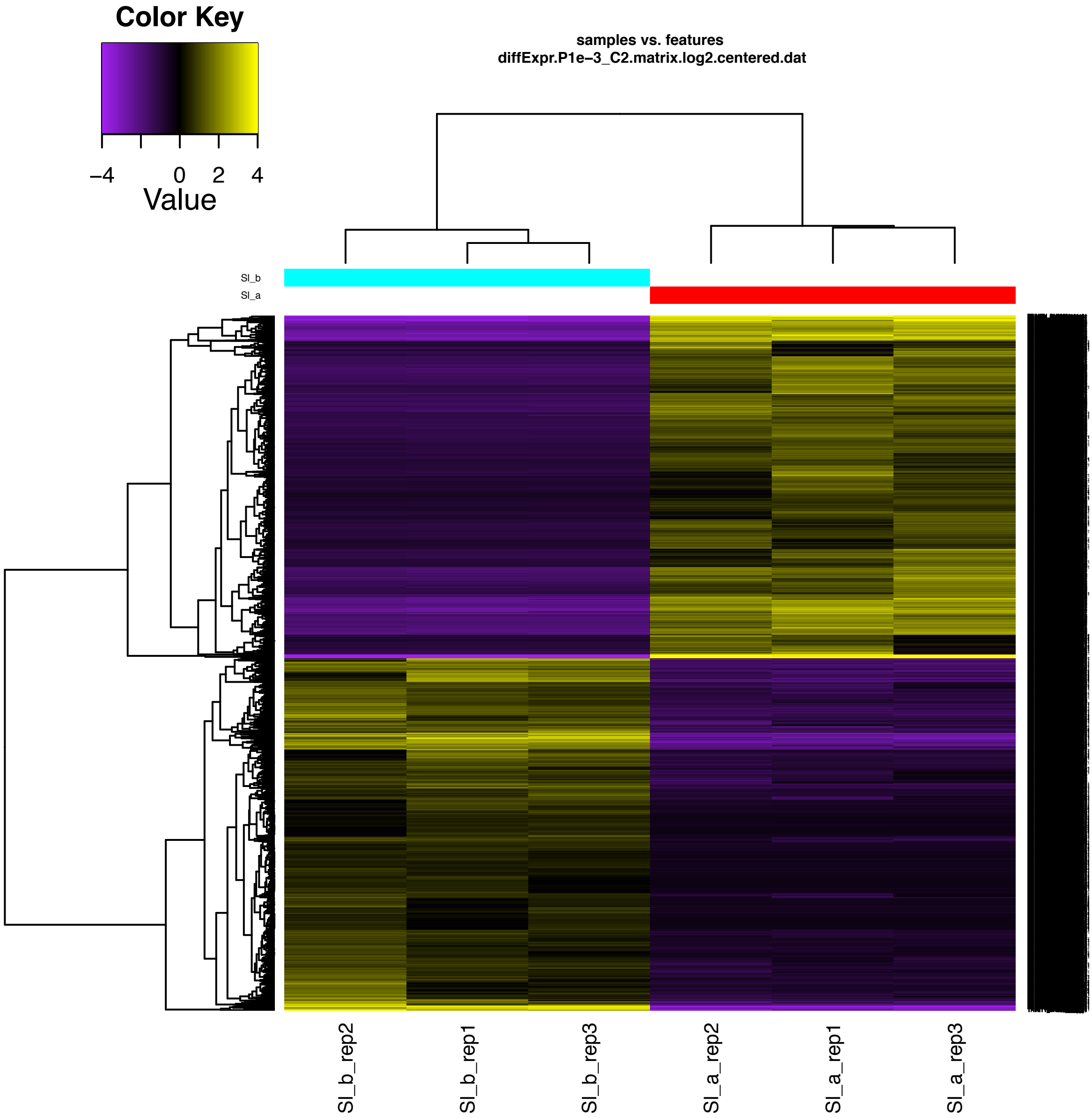
