## supplementary figure S4 for "Smelling in the dark: phylogenomic insights on the chemosensory system of a subterranean beetle"

**Supplementary figure S4.** Alignment of all isoforms of IR25a candidates for *S. longicornis.* In bold, the longest isoforms used to infer the phylogenies.

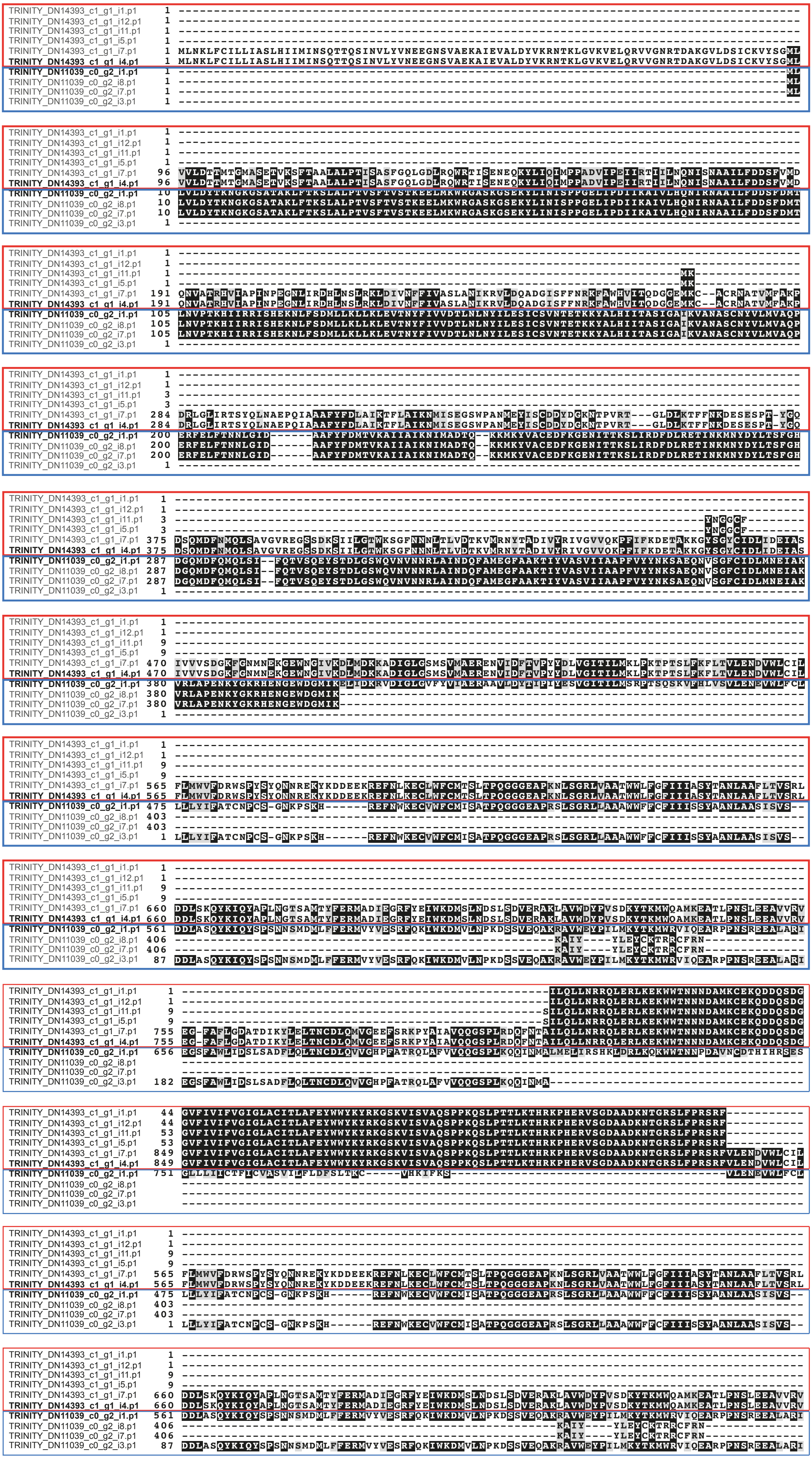
